# Dynamic population stage-structure stabilizes complex ecological communities

**DOI:** 10.1101/2020.06.27.174755

**Authors:** André M. de Roos

## Abstract

Natural ecological communities are diverse, complex and often surprisingly stable, but the mechanisms underlying their stability remain a theoretical enigma^1-5^. Interactions such as competition and predation presumably structure communities^6^, yet theory predicts that complex communities are only stable when species growth rates are mostly limited by intraspecific self-regulation rather than by interactions with resources, competitors and predators^3,5,7^. Current theory, however, only considers the network topology of population-level interactions between species and neglects within-population differences among juvenile and adult individuals. Here, using model simulations, I show that including commonly observed differences in vulnerability to predation and foraging efficiency between juvenile and adult individuals results in up to ten times larger, more complex communities than in simulations without population stage-structure. These diverse communities are stable or fluctuate with limited amplitude, even though in the model only a single basal species is self-regulated and the population-level interaction network is highly connected. Analysis of the species interaction matrix predicts the simulated communities to be unstable but extending the matrix with a population structure subsystem reveals that dynamic changes in population stage-structure completely cancel out this instability. Common differences between juveniles and adults and fluctuations in their relative abundance hence have a decisive influence on the stability of complex natural communities and their vulnerability when environmental conditions change. Thus, community persistence can not be explained by the network of interactions between the constituting species alone.

## Main text

Almost half century ago Robert May concluded that large, complex ecological communities are unlikely to be stable^3,8^, contradicting prevailing ideas that complexity begets stability^1,2^. May analyzed properties of the community or interaction matrix^9^, which captures the per-capita effect of species on each other’s growth rate and mathematically corresponds to the linearization of an ecological model around an equilibrium point. May’s work started the ongoing diversity-stability debate in ecology^4^ and the search for special characteristics of natural communities and constraints on the interactions between their species promoting stability^10,11^. Subsequent studies have used May’s community matrix approach or simulations of community dynamics to reveal the effect on community stability of, among others, weak interactions between species^12^, adaptive foraging^13^, allometric scaling of interaction strength^14^, omnivorous^15^ and mutualistic interactions^16^.

Modelling studies on community stability mainly focus on different aspects of the food web, the ecological network of interspecific interactions. But all of them necessarily involve assumptions about the direct, negative effect of the density of species on their own growth rate. Such self-effects or self-regulation might come about through, for example, cannibalism, interference competition for available nest sites or fighting among predators while foraging. To what extent self-regulation occurs in natural populations is debated^15,17-19^ and empirical assessment of its strength is challenging. Community models, however, require self-regulation for community stability^5,17,18^. To what extent mechanisms such as weak interspecific interactions are stabilizing is even measured in terms of how much self-regulation is required for community stability^12,20^. A recent review concluded that for community stability to occur “at least half—and possibly more than 90%—of species must be subject to self-regulation to a substantial degree”^5^, even though clear empirical evidence for it is lacking.

According to existing theory, stability of ecological communities therefore hinges on negative interactions between individuals of the same species, more than on their interactions with resources, competitors or predators, which contrasts with the idea that species interactions, such as competition and predation, structure ecological communities^6,21^. Studies of ecological networks and food webs have focused on topological characteristics of the network of species interactions in a community, the nature of these interactions and their strength. The species or population is adopted as the basic unit of measurement or modelling and within-population variability between individuals is neglected, even though in real populations no two individuals are the same. The most obvious source of within-population variability results from demographic differences between juvenile and adult individuals as juveniles only grow and mature, while only adults reproduce. Differences between juvenile and adult body size, however, also entail differences in their ecology^22,23^. Because juveniles are smaller they generally feed at lower rates^24^ and are more vulnerable to predation^25,26^. These body size-dependent differences translate into an asymmetry in the ecological interactions of juveniles and adults, which has been shown to affect the structure and dynamics of ecological communities^27,28^.

To study the impact of juvenile-adult foraging and predation asymmetry on community diversity and stability I randomly constructed replicate model food webs consisting of species with different body weights and compared their dynamics with and without accounting for population structure (see Methods). Only the species with the smallest body weight – the basal species – was assumed to follow self-regulating population dynamics, foraging interactions between all non-basal species were modeled based on the difference in species body weight^29,30^ (see Methods and Extended Data Figure 9). Juveniles and adults of the same species often forage on different resources^31^. Such ontogenetic diet shifts may have major consequences for community structure, for example leading to alternative stable community equilibria^32^. To focus exclusively on differences in foraging rate and predation mortality of juveniles and adults and to allow comparison between models with and without population structure, I excluded the possibility of ontogenetic niche shifts and assumed juveniles and adults to forage at possibly different rates on the same range of prey species with the same preferences and thus have overlapping diets. Juvenile and adult dynamics were modeled using a juvenile-adult structured model^33^ in terms of numerical abundances that explicitly accounts for maintenance requirements, which causes maturation and reproduction to halt at low food availabilities (see Methods). Asymmetry in resource foraging is represented phenomenologically by a foraging asymmetry factor *q*, ranging between 0 and 2, with juvenile and adult resource ingestion rate taken proportional to *q* and (2-*q*), respectively. For *q*=1 juveniles and adults hence forage at the same rate, such that maturation and reproduction are limited by food to the same extent. Asymmetry in vulnerability to predation is represented analogously by a predation asymmetry factor *ϕ*, also ranging between 0 and 2, with predation mortality of juveniles and adults taken proportional to *ϕ* and (2-*ϕ*), respectively. For *q*=1 and *ϕ*=1 juveniles and adults hence do not differ in their rates of foraging and predation mortality and the stage-structured food web model is analytically equivalent to a food web model without stage structure (see Supplementary Information, section 1). Community dynamics were simulated until no further species extinctions occurred and the fluctuations in density of the persisting species had stabilized (see Methods).

When juveniles feed at lower rates and are more predator-sensitive than adults (*q*=0.7, *ϕ*=1.8) the structured model results in communities with on average 20 or more non-basal species persisting on the single basal species (Figure 1). Food web simulations with the corresponding unstructured model result in long-term persistence of on average 3-4 non-basal species (Figure 1, Extended Data Figure 1). This increase in community diversity due to juvenile-adult asymmetry is larger at higher system productivity (Extended Data Figure 2). The juvenile-adult asymmetry hence increases the diversity of communities resulting from the food web simulations with at least a factor of 5. Juvenile-adult asymmetry also increases the complexity of the food web that structures the community. Food webs resulting from model simulations without stage-structure are simple, mostly linear food chains, with the majority of species foraging on at most a single prey and vulnerable to a single predator (Figure 1 and Extended Data Figure 3). In contrast, food webs resulting from simulations with juvenile-adult asymmetry are complex with most species foraging on multiple prey species and exposed to predation by multiple consumer species (Figure 1 and Extended Data Figure 3). Diverse and complex communities occur in particular when predation on juveniles is 8-10 times larger than on adults and is less dependent on foraging differences (Extended Data Figure 1; see also Extended Data Figure 6). Judging from the fact that the ratio between body sizes of individual predators and the prey individuals in their gut tends to be an order of magnitude larger than the ratio of average body sizes in the predator and prey populations^30,34^, the asymmetry between predation on juveniles and adults assumed in the simulations seems to be supported by observations.

**Figure 1.**
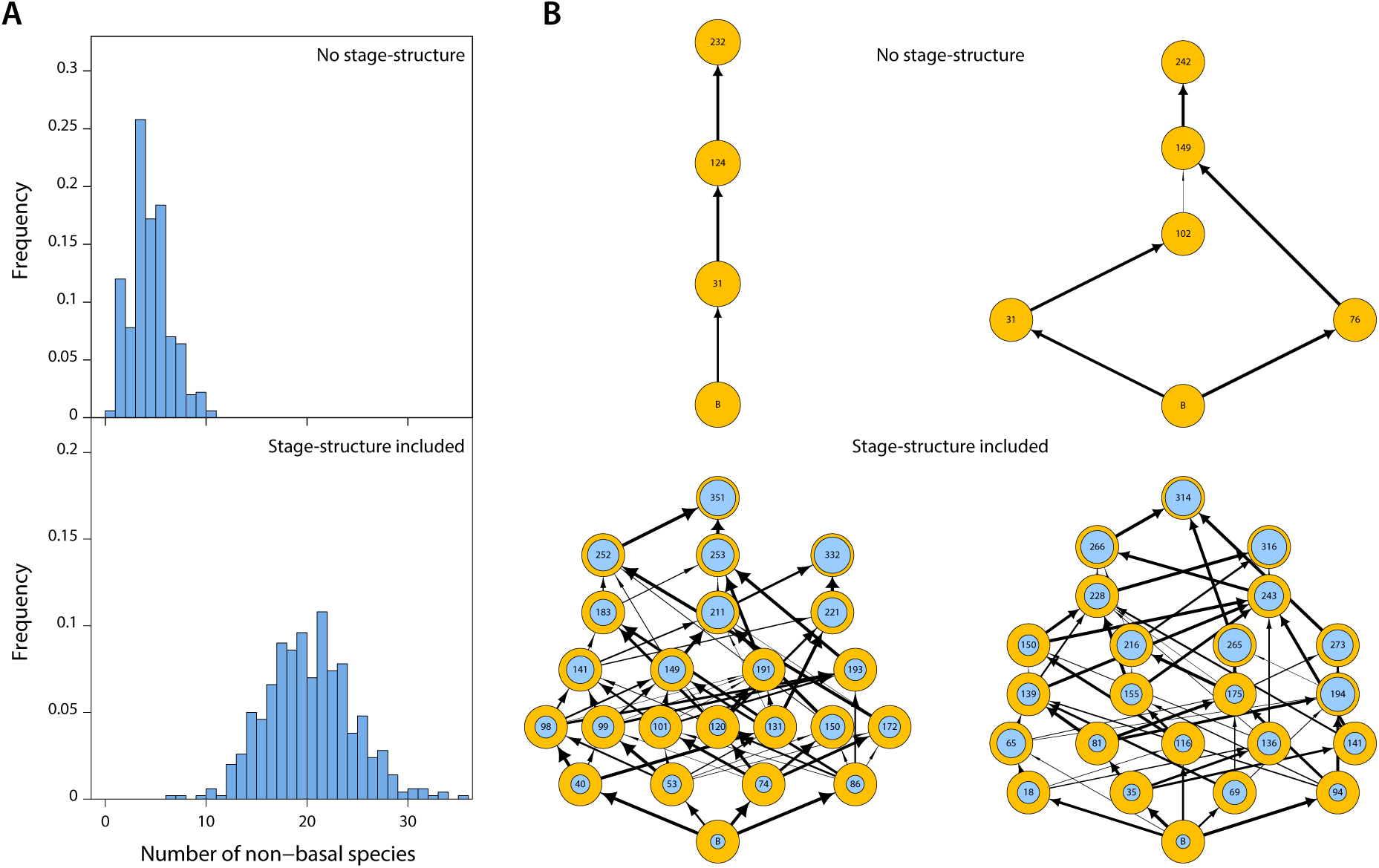
Juvenile-adult stage-structure increases community size and complexity. *A*: Frequency distribution of community sizes (non-basal species only) resulting from 500 replicate food web simulations without (*top*) and with stage-structure and foraging and predation asymmetry between juveniles and adults (*bottom*; *q* = 0.7, *ϕ* = 1.8, see Methods). *B*: Examples of food webs resulting from simulations without (top row) and with stage-structure and foraging and predation asymmetry between juveniles and adults (bottom row; *q* = 0.7, *ϕ* = 1.8). Vertical position indicates trophic level. Inner circles in bottom row indicate the density of juveniles as fraction of total population density. Arrow widths indicate the relative feeding preference (*ψ*_*ik*_, see Methods) of consumers for a particular prey species.

Temporal dynamics of food webs that result from model simulations without stage-structure are characterized by large-amplitude fluctuations in species abundances reminiscent of classical predator-prey or paradox-of-enrichment cycles^35^ (Figure 2). The cycle amplitudes moreover increase with increasing community size especially because minimum species densities during the cycle decrease (Figure 2), ultimately leading to species extinction and reductions in community diversity. In contrast, the complex food webs resulting from the structured model with juvenile-adult asymmetry are either stable or exhibit small-amplitude fluctuations in total species density (Figure 2). When juveniles are inferior resource foragers and more predator-sensitive than adults 22% (115 out of 500 simulations) of the food webs generated by the structured model even approach a stable community equilibrium. Furthermore, if fluctuations in total species density occur, their amplitude is unaffected by community size (Figure 2).

**Figure 2.**
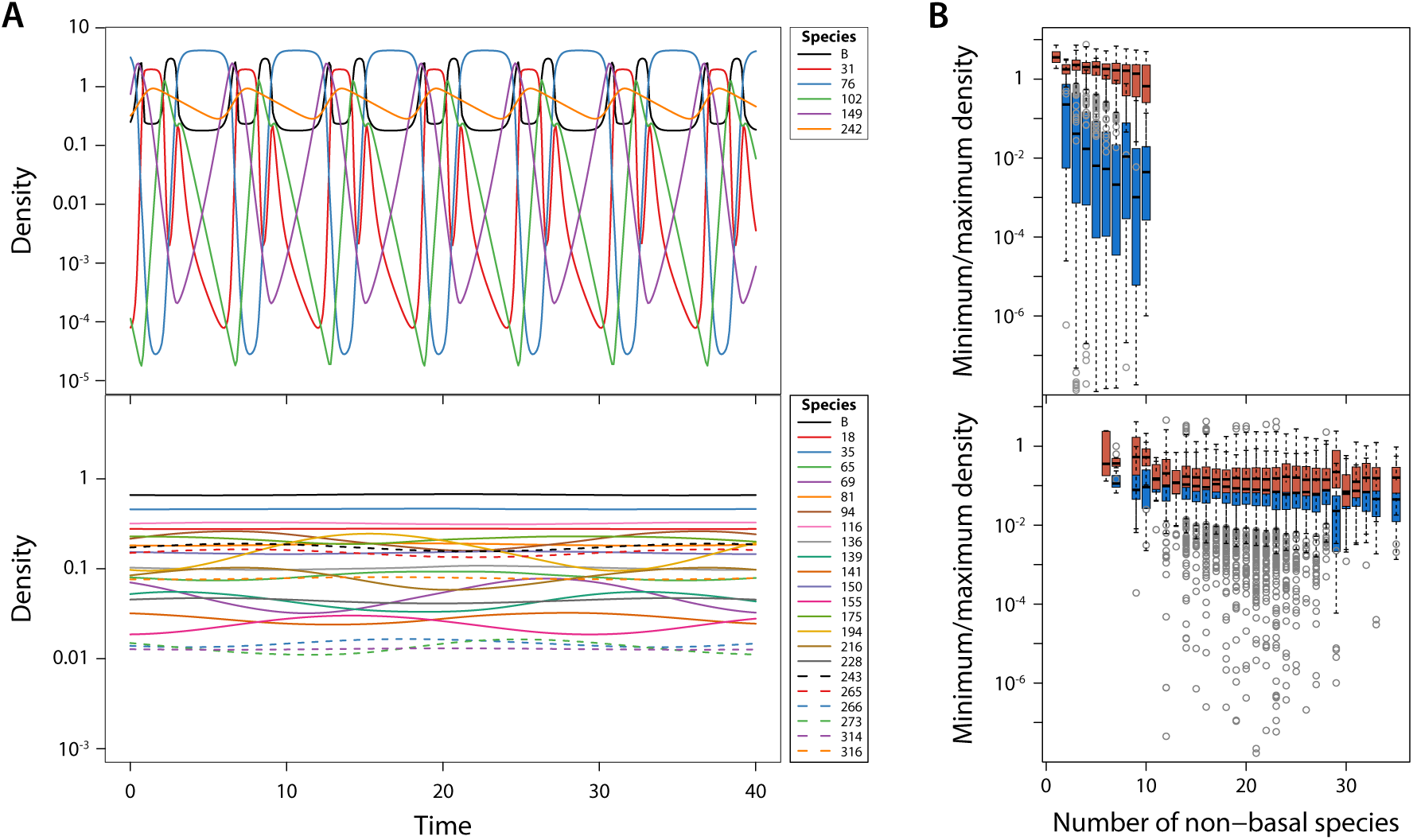
Juvenile-adult stage-structure stabilizes community dynamics. *A*: Examples of dynamics of all species in food web simulations without (top row) and with stage-structure and foraging and predation asymmetry between juveniles and adults (bottom row; *q* = 0.7, *ϕ* = 1.8, see Methods). Corresponding food web structures are shown in Figure 1 (panel B, right column). *B*: Boxplot of minimum (blue bars) and maximum total densities of all populations as a function of community size for all persisting species in 500 replicate food web simulations without (top panel) and with stage-structure and between juveniles and adults (bottom panel; *q* = 0.7, *ϕ* = 1.8).

The stable communities resulting from the structured model with juvenile-adult asymmetry allow for disentangling how food web interactions and population structure together affect community stability. The structured model in terms of juvenile and adult abundances can mathematically be transformed into an equivalent model, in which each species is represented by its total density and the fraction of juveniles in its population. The transformation separates the system of equations into a ‘species-density subsystem’ and a complementary ‘species-structure subsystem’ (see Methods). Community stability is now determined by the stability of each of these two subsystems on their own, and their interactions (see Methods and Supplementary Information section 2). The stability of the species-density subsystem on its own is determined by the standard community matrix, measuring the per-capita effect of species on each other’s growth rate. For all stable communities resulting from the structured model in case of juvenile-adult asymmetry the dominant eigenvalue of this community matrix is positive and large (Figure 3). The community matrix hence predicts these communities to be unstable, which mostly results because only the basal species is regulated by a negative self-effect, whereas non-basal species that experience predation exhibit positive self-effects and top predators have no self-effect (Extended Data Figure 4). The dominant eigenvalue of the Jacobian matrix determining the stability of the coupled subsystems of species density and species structure, however, has a negative real part for all stable communities (Figure 3) mostly because the dominant eigenvalue of the matrix determining the stability of the species-structure subsystem on its own has a negative real part (results not shown). The large differences between the dominant eigenvalues of the Jacobian and community matrices indicate that the dynamic nature of the fraction of juveniles of the species is key to community stability.

**Figure 3.**
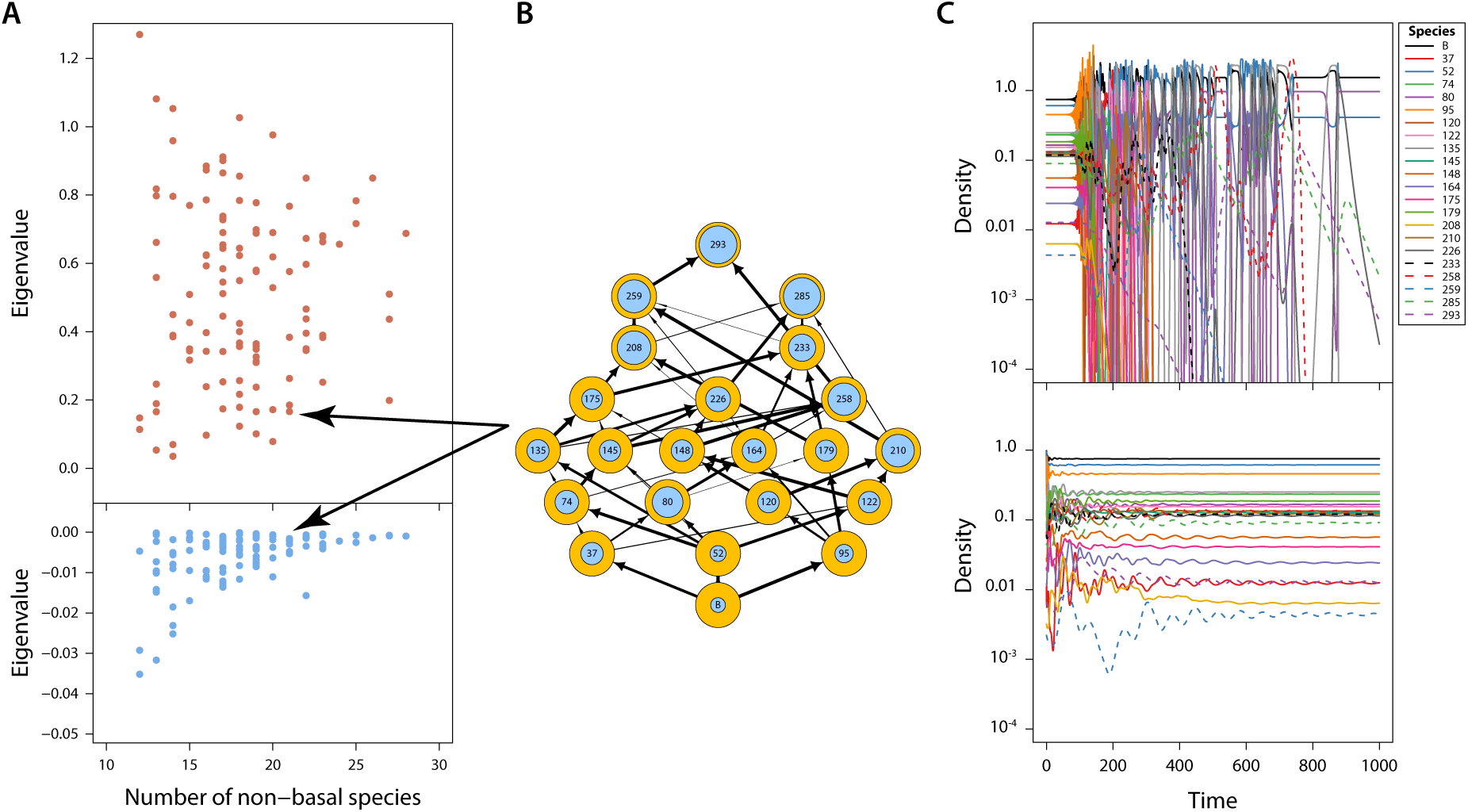
Adaptive stage-structure stabilizes community dynamics. *A*: Real part of the dominant eigenvalue of the community matrix (*top*) and the Jacobian matrix (*bottom*) as a function of community size for all communities that on the basis of simulations with the stage-structured model and foraging and predation asymmetry between juveniles and adults are judged stable (*q* = 0.7, *ϕ* = 1.8; see Methods and Supplementary Information, section 2). *B*: Example of a stable community with 21 non-basal species. *C*: Dynamics of the community shown in *B* with a constant juvenile-adult density ratio equal to its equilibrium value for each species and initial densities equal to their equilibrium densities (*top*) and with dynamic juvenile-adult stage-structure following a disturbance event that reduces the densities of all species in the community by 50% (*bottom*; see Methods).

Simulations of community dynamics starting from these equilibrium community states confirm the stabilizing impact of the population structure and how it increases the resilience of the community: Even after a disturbance event that reduces the density of all species by 50% the complete model involving the coupled species-density and species-structure subsystems predicts a rapid return to the stable community equilibrium (Figure 3). The reduction in density at most results in the extinction of a few species (Extended Data Figure 5). In contrast, when starting in the undisturbed community equilibrium and simulating dynamics using only the species-density subsystem with for each species the fraction of juveniles constant in time and equal to its equilibrium value, species densities only briefly stay close to their equilibrium values and start to fluctuate wildly afterwards (Figure 3). As a result, most species eventually go extinct and the community ends up being of similar size as the communities predicted by the food web model without population structure (Extended Data Figure 5). The dynamics of the juvenile fraction in each of the species dampens density fluctuations in the community in various ways. For example, a perturbation increasing the reproduction of one particular species would have more limited effects on its prey species and be more quickly quenched by its predators, if juveniles forage at lower rates and are more vulnerable to predation, than in the absence of any population structure. Furthermore, a dynamic population stage-structure also buffers against fluctuations in predation pressure, as increases in predation will primarily affect the juveniles that are limited most by food availability. Higher mortality under these conditions has been shown to relax possible bottlenecks in juvenile maturation and to increase the efficiency with which resources are used for population growth as opposed to being used for somatic maintenance^33,36^.

The presented results are robust to changes in the population stage-structure as well as the model describing dynamics of each of the species. Similar results regarding community diversity, complexity and dynamics are obtained under even wider parameter ranges when populations are represented by the biomass densities in 3 life history stages (small juveniles, larger immatures and adults) as opposed to numerical abundances of juveniles and adults only and the dynamics of each population is modelled using a stage-structured biomass model that approximates the dynamics of a complete population body size-distribution^37^ (Extended Data Figures 6-8, Supplementary Information section 3). The communities resulting from this more detailed model tend to be even larger with on average 25-30 species coexisting on a single basal species.

Introducing population stage-structure and its dynamics can be viewed as extending the food web to a multilayer ecological network of species interactions^38^ by including complementary juvenile and adult layers. However, nodes (species) in the juvenile and adult layer of the network always persist or perish together, as neither juveniles nor adults can persist on their own. Furthermore, juvenile density only increases through adult reproduction and adult density only through juvenile maturation. This dynamic dependence of each of the life history stages on its complementary stage sets population stage structure apart from other types of within-population heterogeneity, for example, arising as a result of trait variability in a population, as the dynamics of subgroups with different traits are not necessarily dependent on each other. The presented results contrast with earlier studies of the impact of stage-structure on food web dynamics^39^, because the model for the juvenile-adult stage dynamics more realistically accounts for basic maintenance costs of individuals, causing maturation and reproduction to halt at low food availability when food intake is barely enough to cover these costs.

In conclusion, the topology of the interaction network between species in a community may only provide limited insight into the mechanisms stabilizing complex communities and may even suggest necessary conditions for stability, such as ubiquitous self-regulation, that turn out to be redundant once the dynamics of population stage-structure is taken into account. The mechanisms originating from a dynamic population structure simply overrule the destabilizing effects of the species interaction network. In a realistic and natural way population stage-structure dynamics thus breaks up the constraints on community complexity that were originally identified by May^3,8^. Furthermore, accounting for differences between juveniles and adults in community modelling opens up a direction in food web ecology with new questions. For example, about the implications of the commonly observed ontogenetic niche and diet shifts^31^ for the complexity and stability of communities and about the extent to which population stage-structure promotes the resilience of communities exposed to stress or disturbances. If we are to model the impact of environmental change on complex ecological communities, we need models that can fully capture the diversity, complexity, stability and vulnerability of these systems – this study represents a major advance on current approaches that only consider species-level interaction networks.

## Supplementary Information

All data files and scripts used to generate the figures can be assessed by installing the R package “StructuredFoodweb” using the command devtools::install_bitbucket(“amderoos/structuredfoodweb”).

## Acknowledgements

I want to thank Bernd Blasius, Jennifer Dunne, Jacopo Grilli, Bob Holt, Kevin Lafferty, Ben Martin, Lennart Persson and Peter de Ruiter for stimulating discussions and their comments on the manuscripts and the Santa Fe Institute, Santa Fe (NM), USA, for the hospitality while carrying out this research.

## Author Contributions

AMdR conceived and carried out this study.

## Extended Data Figures

**Extended Data Figure 1.**
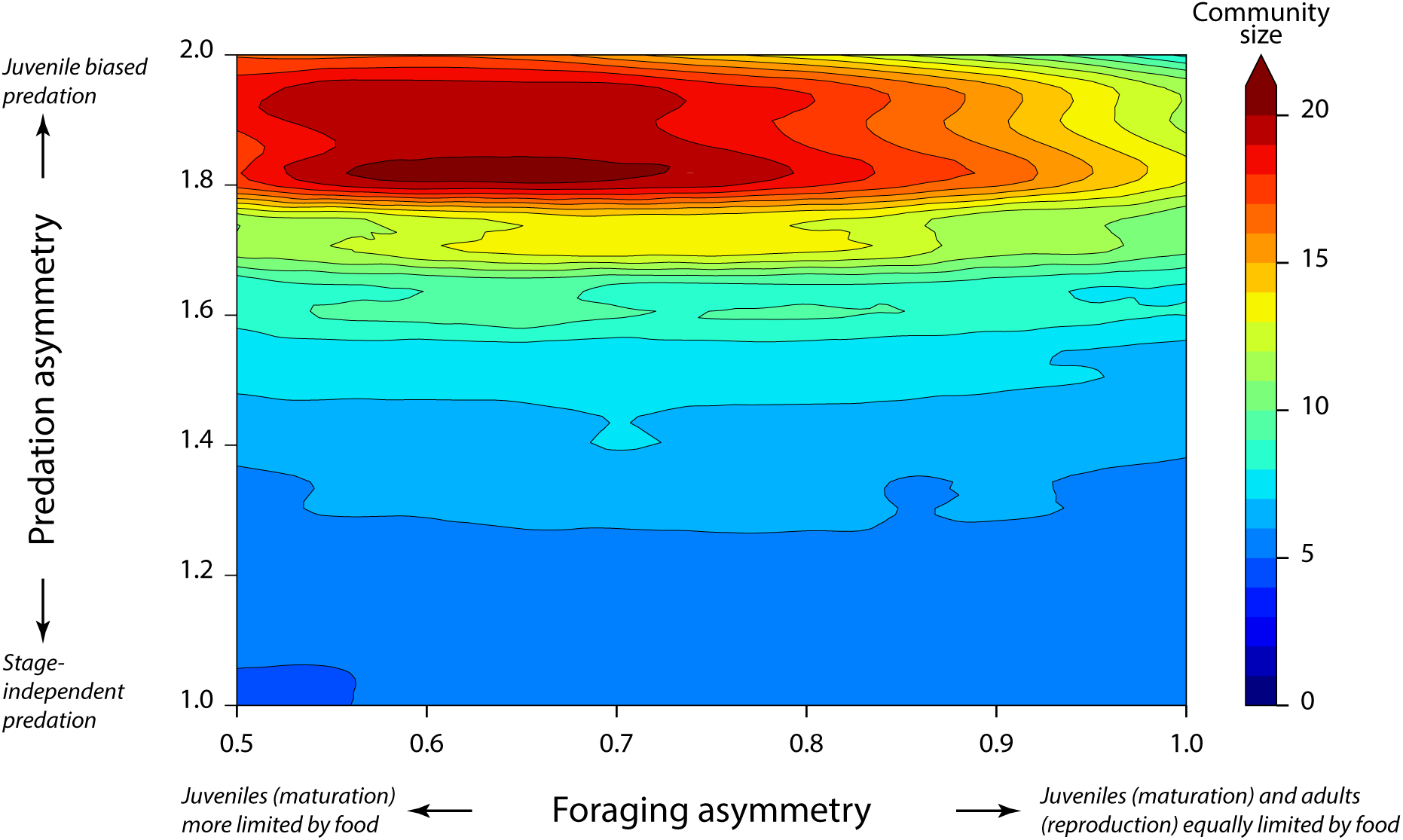
Juvenile-adult asymmetry increases community diversity. Mean community size (non-basal species only) of 500 replicate food web simulations with juvenile-adult stage-structure for different values of foraging (*q*) and predation (*ϕ*) asymmetry between juveniles and adults. Larger communities result when predation is stronger on juveniles than on adults and maturation is more limited by food availability than reproduction.

**Extended Data Figure 2.**
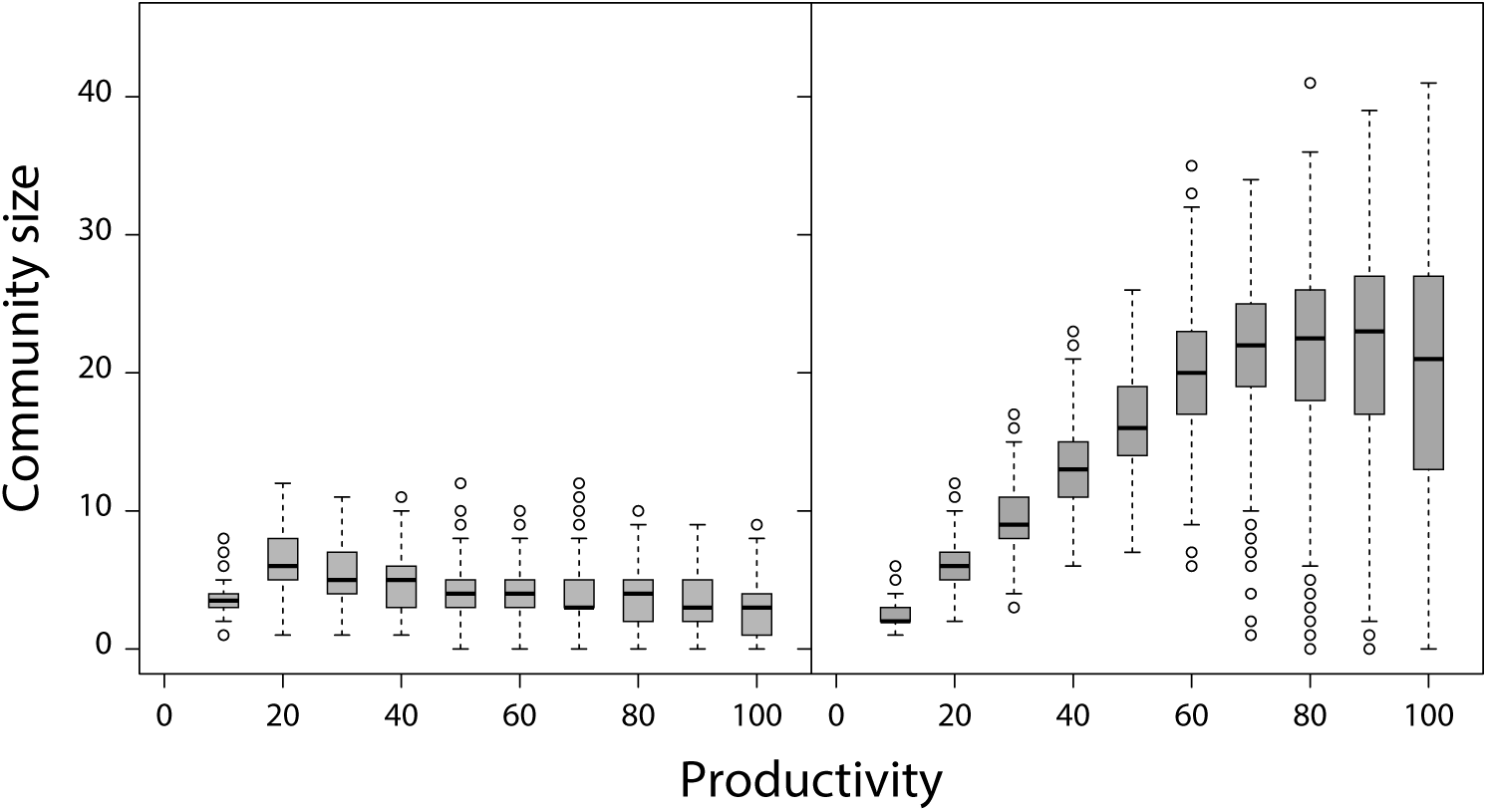
Juvenile-adult asymmetry increases community diversity at all productivities: Boxplot of community sizes at different levels of system productivity *P* resulting from 500 replicate food web simulations without (*left*) and with stage-structure and foraging and predation asymmetry between juveniles and adults (*right*; *q* = 0.7, *ϕ* = 1.8, see Methods).

**Extended Data Figure 3.**
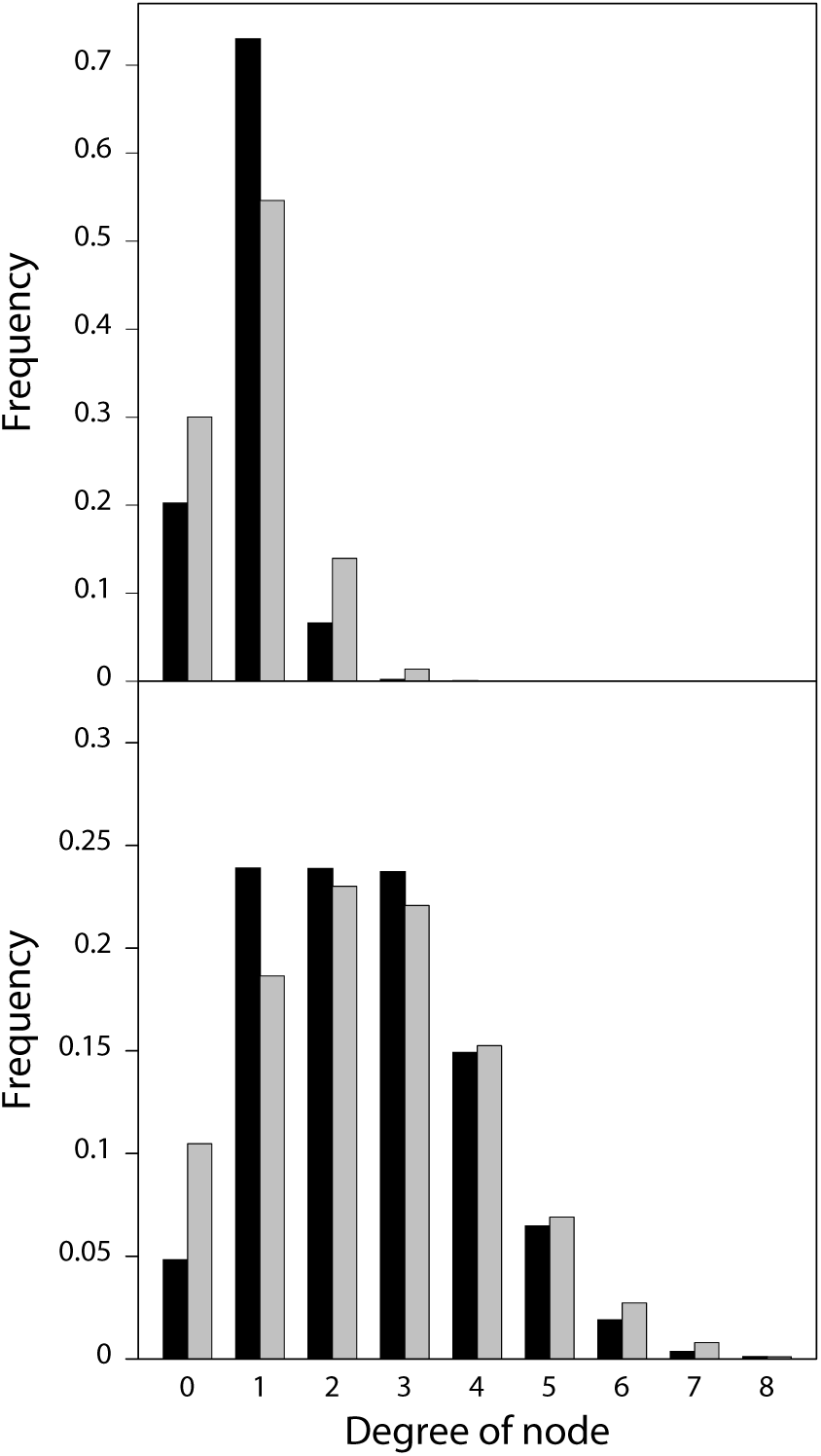
Juvenile-adult asymmetry increases food web connectivity. Number of prey species (black bars; incoming network node links) and predators (grey bars; outgoing network node links) for all species in food webs resulting from 500 replicate simulations without (top panel) and with stage-structure and foraging and predation asymmetry between juveniles and adults (bottom panel; *q* = 0.7, *ϕ* = 1.8).

**Extended Data Figure 4.**
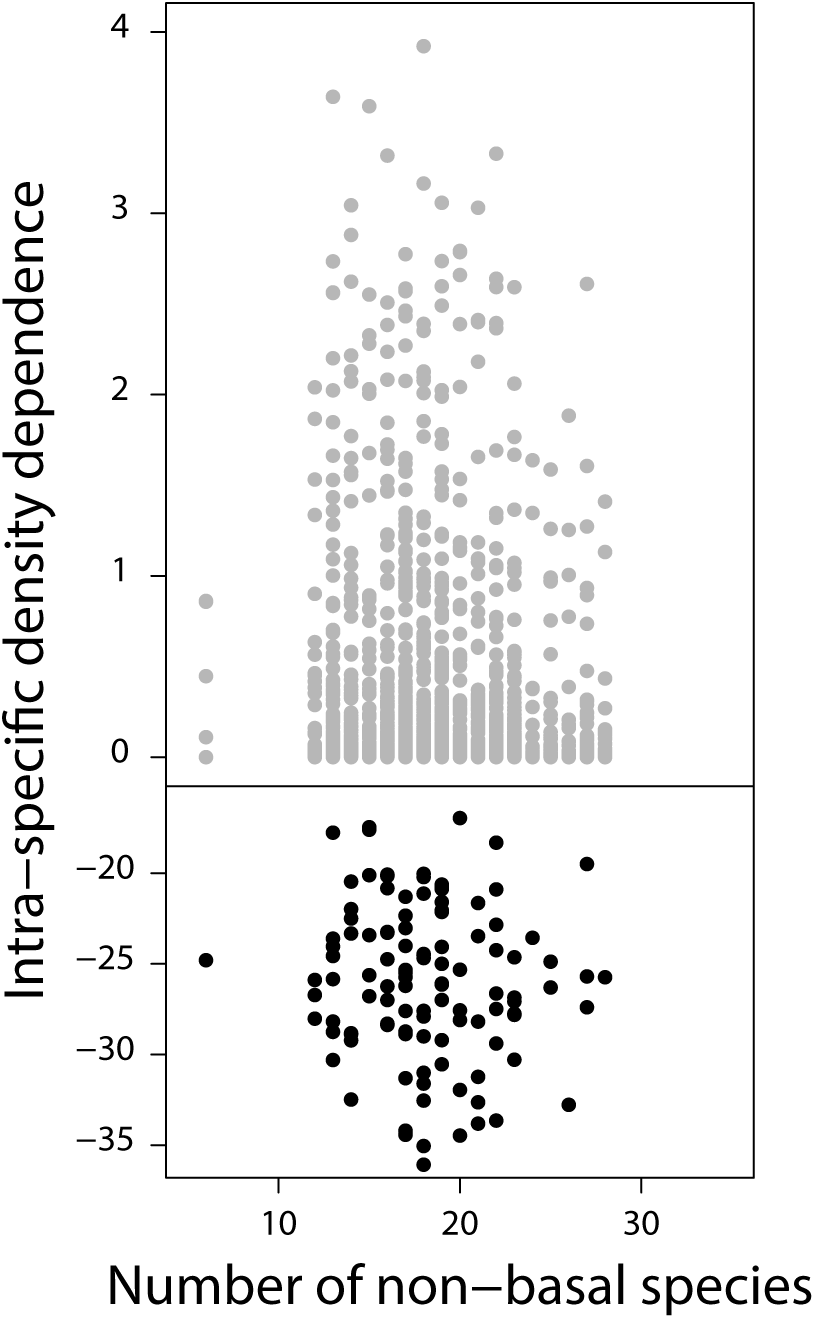
Juvenile-adult asymmetry stabilizes community dynamics without self-regulation. Strength of intra-specific density dependence for basal (*bottom*) and all non-basal species (*top*) in stable communities resulting from food web simulations with the stage-structured model and foraging and predation asymmetry between juveniles and adults (*q* = 0.7, *ϕ* = 1.8). Intra-specific density dependence is assessed with the diagonal elements of the community matrix, which measures for basal and non-basal species the negative and positive effect, respectively, of total species abundance on its own rate of change (see Methods).

**Extended Data Figure 5.**
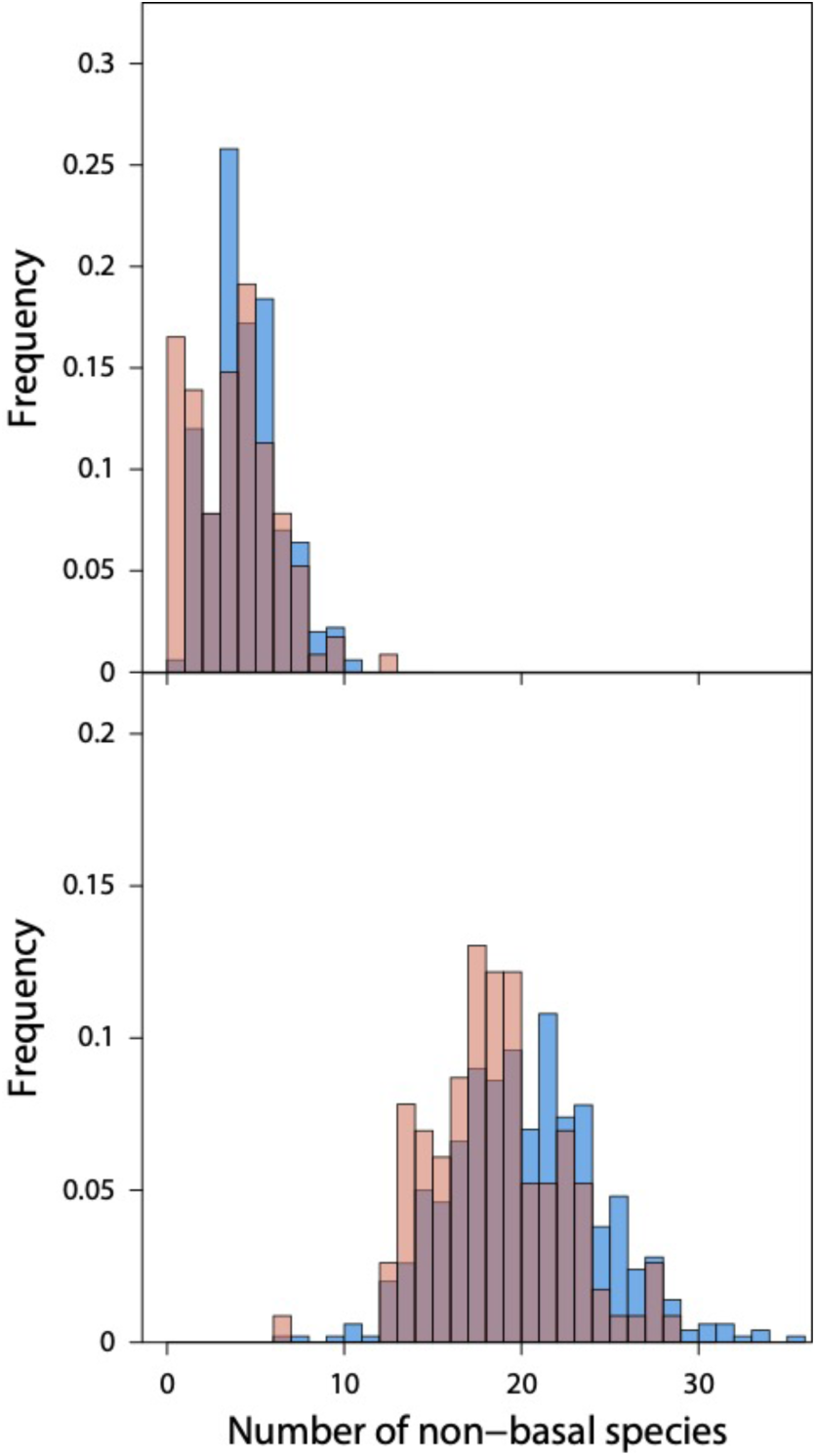
Dynamic juvenile-adult ratio enforces complex community stability. Frequency distribution of community sizes (non-basal species only; red bars) resulting from simulations of dynamics for all stable communities generated by the stage-structured model in case of foraging and predation asymmetry between juveniles and adults (*q* = 0.7, *ϕ* = 1.8; see Methods and Supplementary Information, section 2). Top panel shows results of the species-density subsystem on its own with the juvenile-adult consumer ratio for each species constant and equal to its equilibrium value when initial species densities are identical to their equilibrium values. Bottom panel shows results of the coupled species-density and species-structure subsystem when initial densities for each species equal 50% of their equilibrium densities. For reference, top and bottom panel also show the frequency distribution of community sizes (non-basal species only; blue bars) resulting from 500 replicate food web simulations without and with stage-structure and foraging and predation asymmetry between juveniles and adults, respectively, that are also presented in Figure 1 in the main text.

**Extended Data Figure 6.**
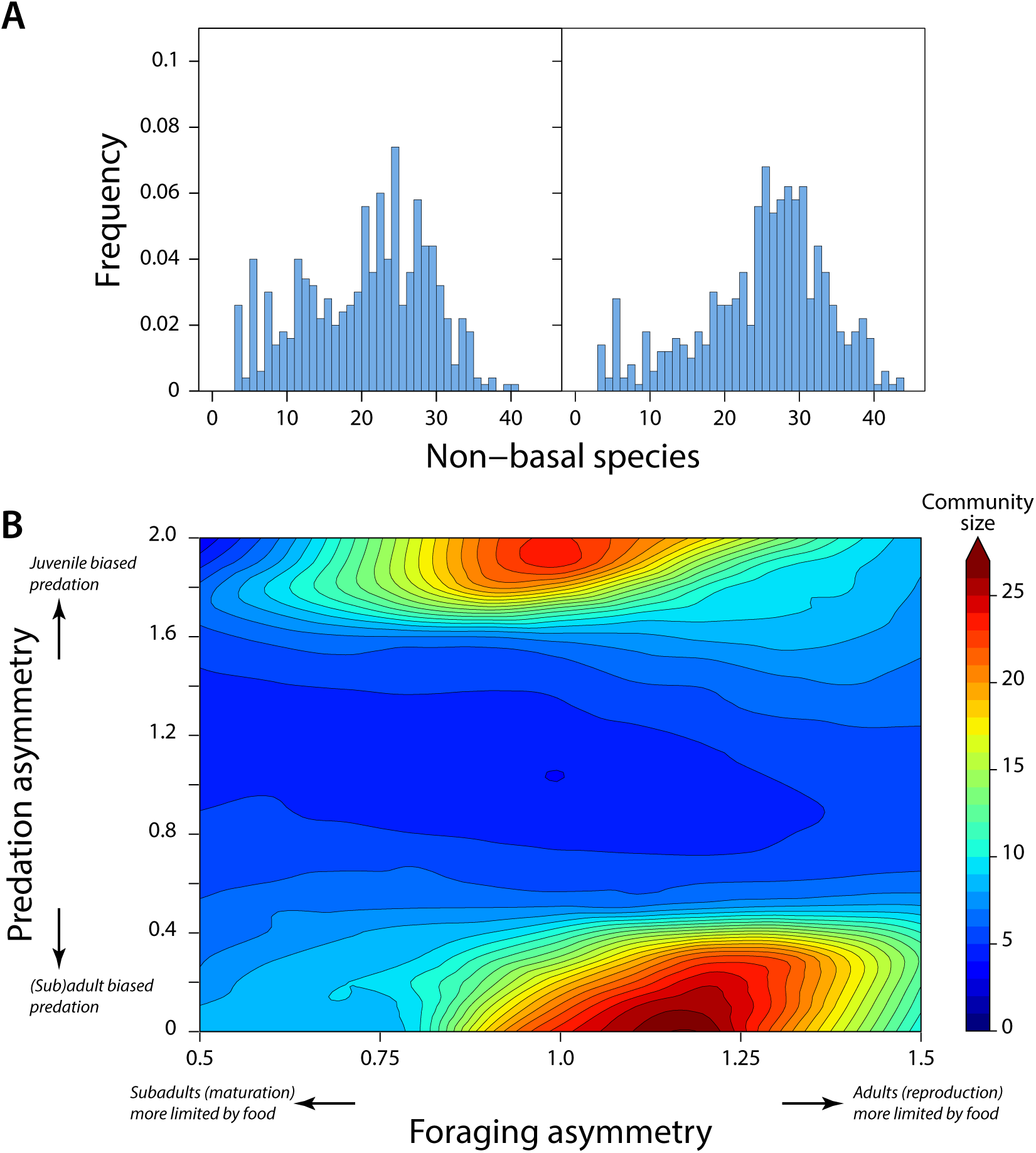
Juvenile-adult asymmetry in biomass dynamics increases community diversity. *A*: Frequency distribution of community sizes (non-basal species only) resulting from 500 replicate food web simulations using the 3-stage biomass model including juveniles, subadults and adults (see Supplementary Information section 3) when juveniles are most vulnerable to predation and subadults are limited by food availability most (left panel; *q* = 0.9, *ϕ* = 1.8) and when subadults and adults are more vulnerable to predation and juveniles are limited by food availability most (right panel; *q* = 1.2, *ϕ* = 0.2). *B*: Mean community size (non-basal species only) of 500 replicate food web simulations using the 3-stage biomass model including juveniles, subadults and adults for different values of foraging (*q*) and predation (*ϕ*) asymmetry.

**Extended Data Figure 7.**
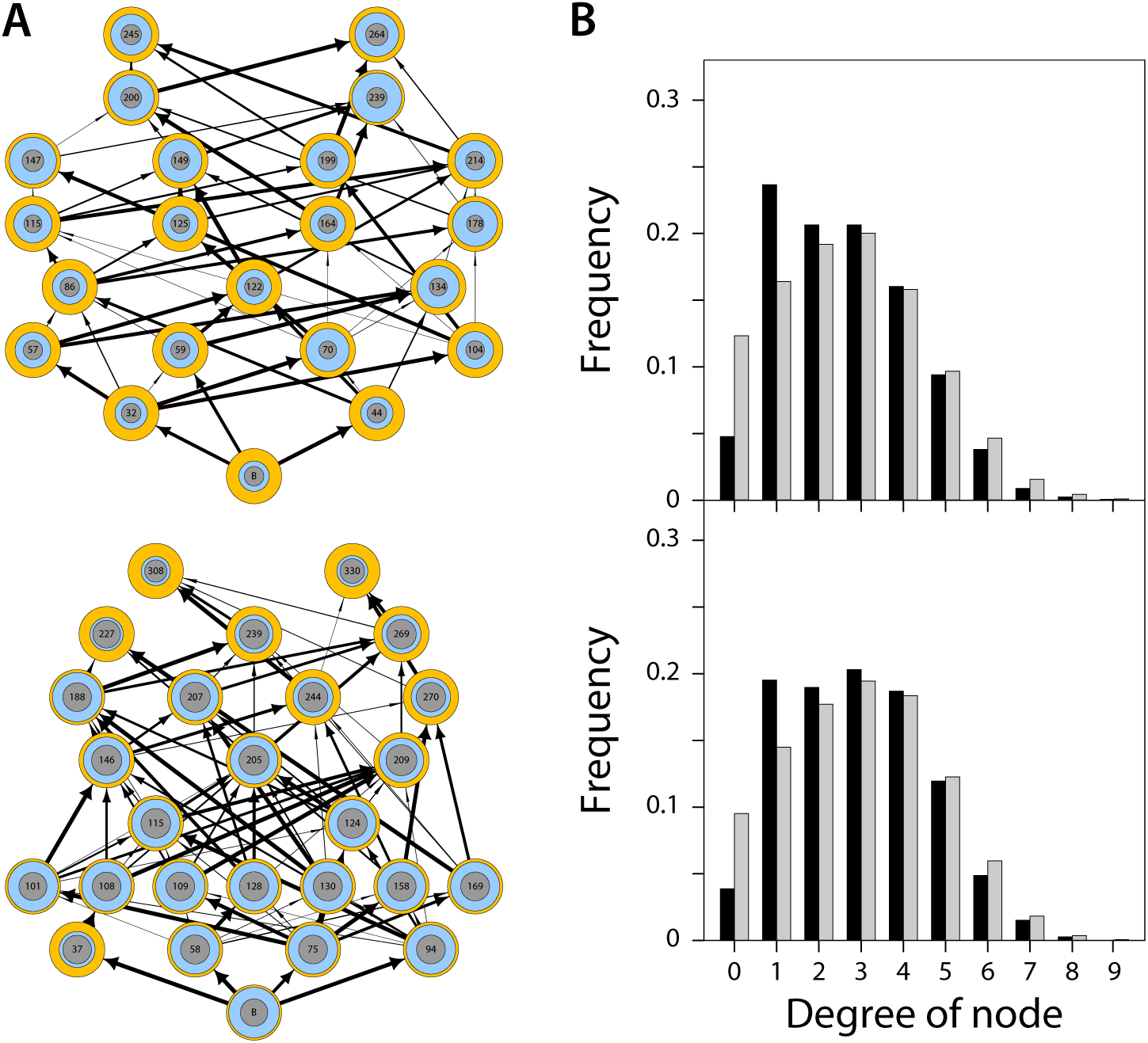
Juvenile-adult asymmetry in biomass dynamics increases food web complexity. *A*: Examples of food webs resulting from simulations using the 3-stage biomass model including juveniles, subadults and adults (see Supplementary Information section 3) when juveniles are more vulnerable to predation and subadults are limited by food availability most (top panel; *q* = 0.9, *ϕ* = 1.8) and when subadults and adults are more vulnerable to predation and juveniles are limited by food availability most (bottom panel; *q* = 1.2, *ϕ* = 0.2). Vertical position indicates trophic level. Inner circles indicate the biomass fraction of juveniles (grey) and total immatures (blue) in the population. Arrow widths indicate the relative feeding preference (*ψ*_*ik*_, see Methods) of consumers for a particular prey species. *B*: Number of prey species (black bars; incoming network node links) and predators (grey bars; outgoing network node links) for all species in food webs resulting from 500 replicate simulations using the 3-stage biomass model including juveniles, subadults and adults when juveniles are more vulnerable to predation and subadults are limited by food availability most (top panel; *q* = 0.9, *ϕ* = 1.8) and when subadults and adults are more vulnerable to predation and juveniles are limited by food availability most (bottom panel; *q* = 1.2, *ϕ* = 0.2).

**Extended Data Figure 8.**
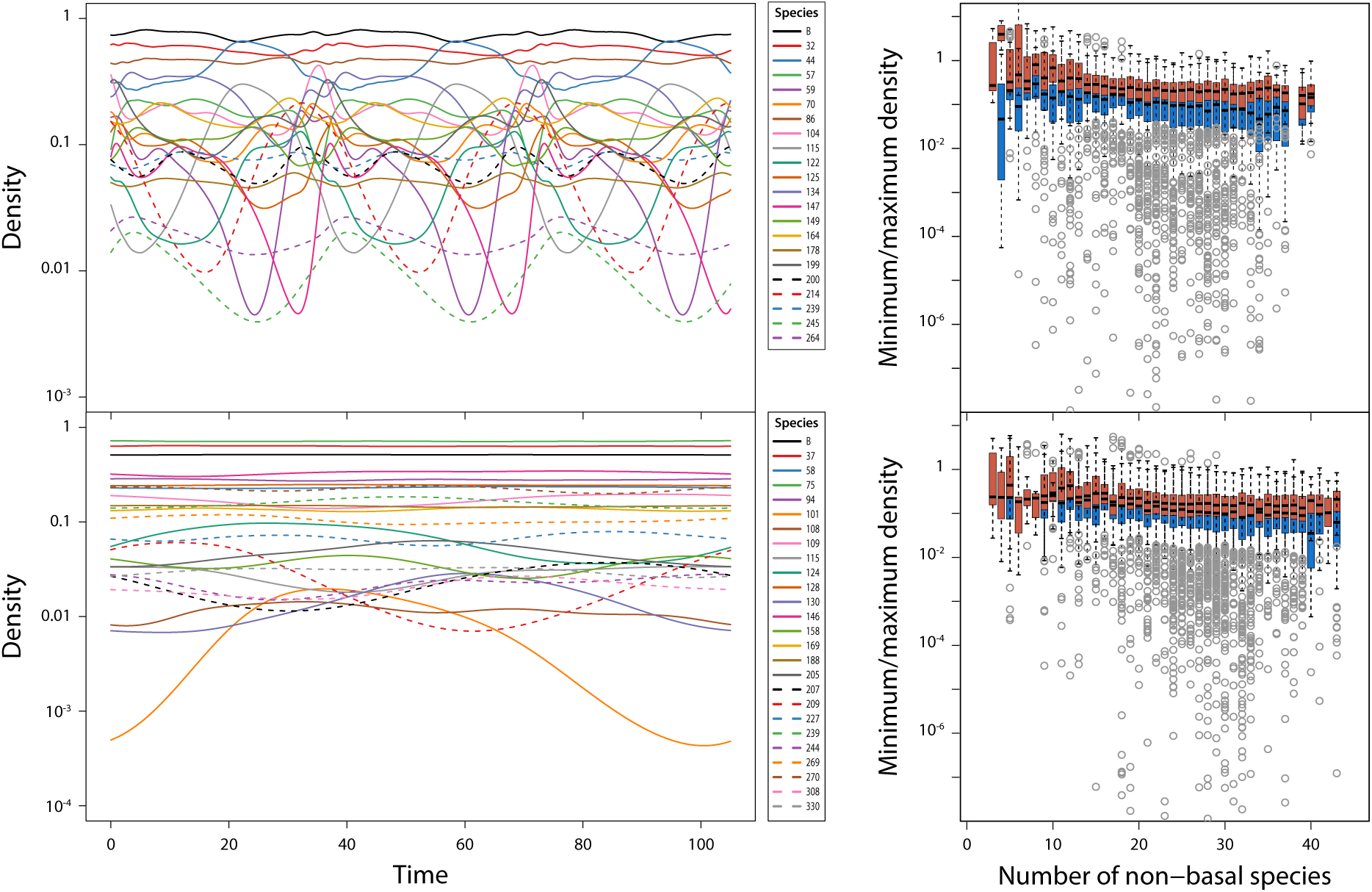
Juvenile-adult asymmetry in biomass dynamics stabilizes community dynamics. *A*: Examples of total biomass dynamics of all species in food web simulations using the 3-stage biomass model including juveniles, subadults and adults (see Supplementary Information section 3) when juveniles are more vulnerable to predation and subadults are limited by food availability most (top panel; *q* = 0.9, *ϕ* = 1.8) and when subadults and adults are more vulnerable to predation and juveniles are limited by food availability most (bottom panel; *q* = 1.2, *ϕ* = 0.2). Corresponding food web structures are shown in Extended Data Figure 7. *B*: Boxplot of minimum (blue bars) and maximum total biomass densities as a function of community size for all persisting species in 500 replicate food web simulations using the 3-stage biomass model including juveniles, subadults and adults (see Supplementary Information section 3) when juveniles are more vulnerable to predation and subadults are limited by food availability most (top panel; *q* = 0.9, *ϕ* = 1.8) and when subadults and adults are more vulnerable to predation and juveniles are limited by food availability most (bottom panel; *q* = 1.2, *ϕ* = 0.2).

**Extended Data Figure 9.**
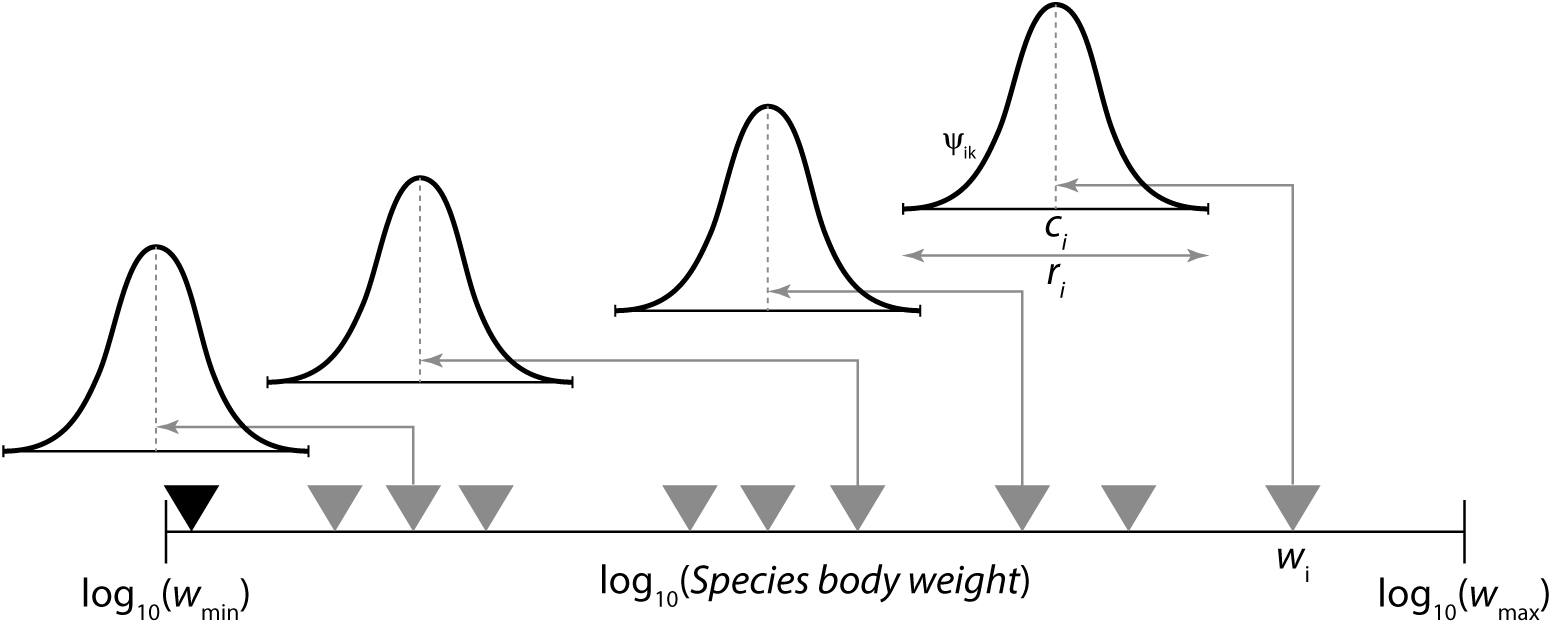
Construction of the prey-predator mass ratio food web model. Species are randomly assigned niche values *n*_*i*_ in the range [0,1]. Niche values are related to body size *w*_*i*_ following 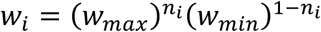 with minimum (*w*_*min*_) and maximum body size (*w*_*max*_) equal to 10^−8^ and 10^4^ gram, respectively. The center *c*_*i*_ of the feeding niche of consumer species is uniformly distributed between *n*_*i*_ −2.5/^10^log(*w*_*max*_/*w*_*min*_) and *n*_*i*_ −0.5/^10^log(*w*_*max*_/*w*_*min*_), yielding median prey-predator body size ratio between 10^−2.5^ and 10^−0.5^. The feeding niche width *r*_*i*_ equals 1/^10^log(*w*_*max*_/*w*_*min*_). Consumer species *i* feeds on all prey species *k* with body sizes between 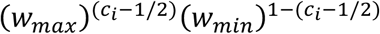 and 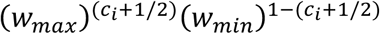 at a relative feeding rate *ψ*_*ik*_ following a hump-shaped distribution of prey body size (see Methods).

## Methods

### Food web construction

Model food webs are constructed by assigning each of the initial *N* = 500 species a random niche value *n*_*i*_ ranging between 0 and 1. Niche values are related to species body size *w*_*i*_ following 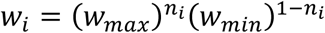 with a minimum (*w*_*min*_) and maximum species body size (*w*_*max*_) equal to 10^−8^ and 10^4^ gram, respectively (see Extended Data Figure 9). To represent documented ratios between prey and predator body sizes^29,30^ more faithfully than in the niche model^40^ the center *c*_*i*_ of the feeding niche of consumer species *i* is uniformly distributed between *n*_*i*_ - 2.5/^10^log(*w*_*max*_/*w*_*min*_) and *n*_*i*_ - 0.5/^10^log(*w*_*max*_/*w*_*min*_), resulting in a median prey-predator body size ratio ranging between 10^−2.5^ and 10^−0.5^. The width *r*_*i*_ of the feeding niche equals 1/^10^log(*w*_*max*_/*w*_*min*_), such that consumer species *i* feeds on prey species with body sizes ranging between 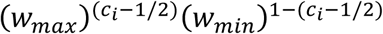 and 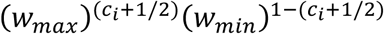. The relative feeding rate *ψ*_*ik*_ of consumer species *i* on prey species *k* in its feeding niche follows a piece-wise continuous hump-shaped distribution with finite range (Bates distribution of order 3):

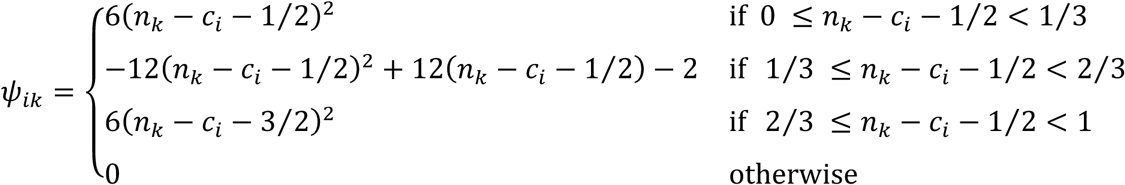

### Food web dynamics without stage structure

Species are ranked according to their niche value (i.e. body size) and their numerical abundances are indicated with *C*_*i*_. The basal species (with index 1) is assumed to forage on its own exclusive resource *R*, the dynamics of which is described by:

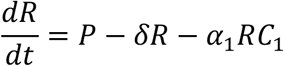

in which *P* is the productivity of the resource and *δ* is its turn-over rate, while *α*_1_ scales the predation pressure of the basal species on the resource. The dynamics of the resource is assumed to be in pseudo-steady state, such that *R* = *P*/(*δ*+*α*_1_C_1_) at all times. The (linear) functional response of the basal species, indicated with *F*_1_, is consequently assumed to equal the pseudo-steady state value of *R*:

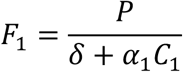

Non-basal species are assumed to forage following a type II functional response on all other species in the community at a relative rate *ψ*_*ik*_ determined by the species body size ratio. The encounter rate of a consumer species with index *i* with all its prey species (indexed with *k*) therefore equals:

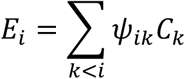

and the value of its functional response *F*_*i*_ (scaled between 0 and 1) equals:

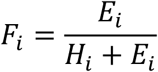

With *H*_*i*_ the consumer’s half-saturation density. Because the prey-predator body size ratio is assumed to be strictly smaller than 1, a consumer species with index *i* can only forage on all species with index *k* < *i*.

The dynamics of all species densities is now described by:

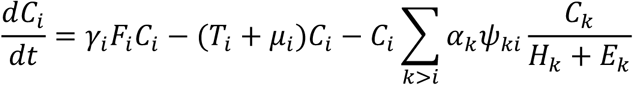

where *E*_*i*_ and *F*_*i*_ represent the value of the food encounter rate and the scaled functional response of species *i*, respectively. The parameter *γ*_*i*_ relates the growth rate of species *i* to its functional response *F*_*i*_, while *α*_*i*_ scales the predation pressure of species *i* on its prey species. The parameters *T*_*i*_ and *μ*_*i*_ represent the population loss rate through somatic maintenance costs and background mortality, respectively. Notice that all species are ordered according to their body size and hence only species with an index *k* > *i* can feed on species *i*.

### Food web dynamics with stage structure

Numerical abundances of juvenile and adult individuals of consumer species *i* are indicated with *J*_*i*_ and *A*_*i*_, respectively. Juveniles and adults are assumed to feed on the same range of prey species, have the same prey preferences and thus overlapping diets. However, juveniles and adults feed at different rates, such that the foraging rates of juveniles and adults of species *i* are proportional to *qF*_*i*_ and (2-*q*)*F*_*i*_, respectively, with proportionality constant *α*_*i*_ and *F*_*i*_ the functional response of species *i*. Juveniles and adults are also assumed to differ in their sensitivity to predation, such that the predation rate of consumers on their juvenile and adult prey is proportional to *ϕM*_*i*_ and (2-*ϕ*)*M*_*i*_, respectively, where *M*_*i*_ represents the predation pressure exerted on species *i* by all of its predators. The parameter *q* and *ϕ* are referred to as the foraging and predation asymmetry between juveniles and adults.

The functional response value for the basal species is defined analogous to the model without stage structure, taking into account the foraging asymmetry between juveniles and adults:

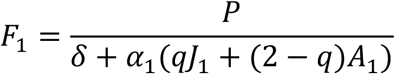

While the encounter rate with prey for non-basal species equals:

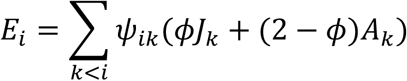

The expression for the functional response of non-basal species is the same as without stage structure:

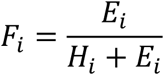

In addition to decreasing through background mortality the numerical abundances of juvenile and adult individuals change through reproduction and maturation. These processes are described by a stage-structured model^33^ that assumes maturation and reproduction to stop when food availability drops below a threshold level and food intake is not sufficient to cover basic maintenance costs. In particular, maturation of juveniles of species *i* depends on its functional response *F*_*i*_ following:

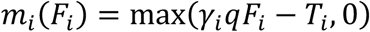

while reproduction by adults follows:

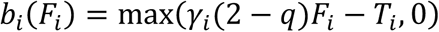

Analogous to the model without stage structure the parameter *γ*_*i*_ relates maturation and reproduction to the food availability, *qF*_*i*_ and (2-*q*)*F*_*i*_, for juveniles and adults, respectively. The maximum functions in the expressions for *m*_*i*_(*F*_*i*_) and *b*_*i*_(*F*_*i*_) ensure that maturation and reproduction halt whenever food availability *F*_*i*_ drops below *T*_*i*_*/*(*qγ*_*i*_) and *T*_*i*_*/*((2-q)*γ*_i_), respectively. The parameter *q* hence determines in a phenomenological manner whether maturation (*q* < 1) or reproduction (*q* > 1) is more limited by food availability. Dynamics of the juvenile-adult structured food web model are described by:

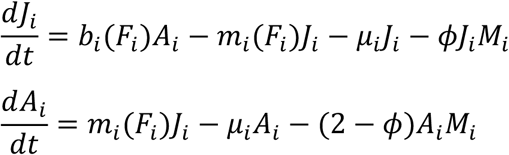

with

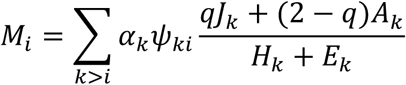

the predation pressure exerted on species *i* by all of its predators.

### Model parameterization

Default parameter values are based on the parameterization as presented by De Roos & Persson^27^ (Box 3.3 and 3.4), except that the time variable and hence all rate parameters have been scaled by a factor of 10 to speed up numerical computations of the model. For each non-basal species, the half-saturation prey density *H*_*i*_ occurring in its functional response *F*_*i*_ was generated independently by sampling from a uniform distribution on the interval [0.5,2.5]. The parameters *α*_*i*_, *γ*_*i*_, *T*_*i*_ and *μ*_*i*_ were assumed to scale with *w*_*i*_^-0.25^. More specifically, for each species *i* the values of these parameters were generated using the following equations:

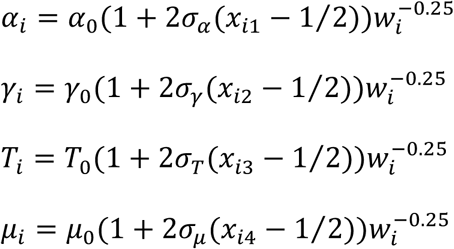

In these expressions *α*_0_=1.0, *γ*_0_=0.6, *T*_0_=0.1 and *μ*_0_=0.015 represent the default mean value of the species-specific parameters^27^. The species-specific parameters *α*_*i*_, *γ*_*i*_, *T*_*i*_ and *μ*_*i*_ were for each species randomly selected from a Bates distribution of degree 3 around these mean values. The Bates distribution is the continuous probability distribution of the mean, *X*, of 3 independent uniformly distributed random variables on the unit interval. Random values from this distribution range between 0 and 1 with mean value of 1/2 and are easily generated by taking the mean of 3 independent samplings from a uniform distribution on the unit interval [0,1]. The quantities *x*_*ij*_ are independent realizations of the random variable *X*, while *σ*_*α*_, *σ*_*γ*_, *σ*_*T*_ and *σ*_*μ*_ represent the one-sided, relative width of the distributions of the species-specific parameters *α*_*i*_, *γ*_*i*_, *T*_*i*_ and *μ*_*i*_, respectively, around their mean values. Default values for these relative widths equal 0.1, such that all species-specific parameters range between 0.9 and 1.1 times their default, mean value and follow hump-shaped distributions within these ranges. Finally, the productivity *P* and turn-over rate *δ* of the exclusive resource of the basal species was taken equal to 60 and 2.0, respectively, in all computations, unless stated otherwise. The two remaining parameters in the model, the foraging asymmetry parameter *q* and the predation asymmetry parameter *ϕ*, were varied between the different computations to assess their effect on community dynamics.

### Numerical simulation procedure

Numerical integrations of the food web with *N* = 500 species were carried out using an adaptive Runge-Kutta (Cash-Karp) method implemented in C. Relative and absolute tolerances during the integration were set to 10^−7^ and10^−13^, respectively. During the first 10000 time units no species were removed from the community, even if they attained very low density. For *t* > 10000 each species, whose total density *J*_*i*_ + *A*_*i*_ dropped below 10^−8^, was removed from the community. This persistence threshold ensures that the product of the relative tolerance (10^−7^) and the lowest species density (10^−8^) is larger than the machine precision (equal to 1.11·10^−16^ according to the IEEE 754-2008 standard in case of double precision). During numerical computations mean and variance as well as the maximum and minimum value of the total species density *J*_*i*_ + *A*_*i*_ were continuously monitored for all species. The values of these measured statistics are reset whenever the community structure changes as one or more species in the community go extinct. Numerical integrations are halted whenever the community structure has not changed for 10^6^ time units and no change has occurred from one time unit to the next in the values of these statistics (mean, minimum, maximum and variance of total species density) for all species in the community.

### Sources of community stability and extent of self-regulation

Through analytical manipulations the model in terms of juvenile abundances *J*_*i*_ and adult abundances *A*_*i*_ can be recast into an equivalent model in terms of total species abundance *C*_*i*_ = *J*_*i*_ + *A*_*i*_ and the fraction of juveniles in a population *Z*_*i*_ = *J*_*i*_/*C*_*i*_. In terms of these alternative model variables the functional response value for the basal species can be written as:

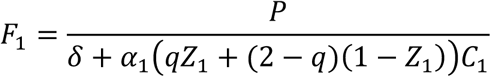

While the encounter rate with prey for non-basal species equals:

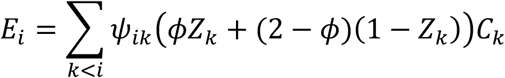

The dynamics of total species density and fraction of juveniles in all populations are then described by:

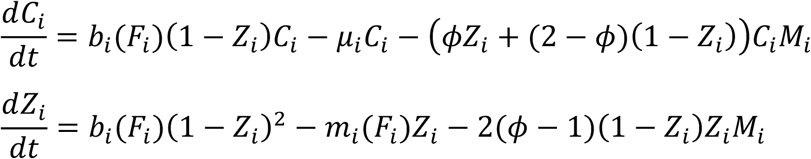

with

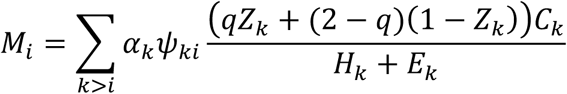

the predation pressure exerted on species *i* by all of its predators.

The resulting system of differential equations can hence be written as:

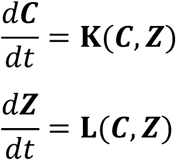

In which ***C*** and ***Z*** indicate vectors of all total species abundances and fractions of juveniles in all populations, respectively. The vector-valued functions **K**(***C, Z***) and **L**(***C, Z***) contain the right-hand side of the differential equations *dC*_*i*_/*dt* for the species-density subsystem and *dZ*_*i*_/*dt* for the species-structure subsystem, respectively. For a system with *m* species the Jacobian matrix **J** of the ODEs above is a 2*m*×2*m* matrix of the form:

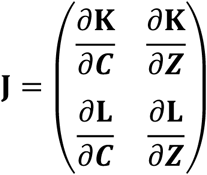

Each of the 4 parts of **J** is a *m*×*m* matrix containing the partial derivatives of the functions **K**(***C, Z***) and **L**(***C, Z***) with respect to the total species densities (*C*_1_, …, *C*_*m*_) and fractions of juveniles (*Z*_1_, …, *Z*_*m*_). Expressions for these partial derivatives are provided in the Supplementary Information (section 2).

All communities resulting from the stage-structured model with asymmetry in feeding and predation between juveniles and adults (*q* = 0.7, *ϕ* = 1.8) for which the minimum and maximum values of the total species density differ less than 10^−6^ from each other for all species are considered stable. For these stable communities the Jacobian matrix **J** is evaluated by substituting for all species the average total abundance and fraction of juveniles observed in the simulation as well as all general and species-specific parameters into the expressions for the elements of **J**. The eigenvalues of the Jacobian matrix are subsequently computed using the routine eigen() in R (Figure 3, left-bottom panel). To evaluate the effect of dynamic changes in population stage-structure (i.e. changes in the juvenile-adult ratio) on the stability of the community equilibrium, these eigenvalues are compared with the eigenvalues of the top-left element of **J**, that is the *m*×*m* matrix ∂**K**/∂***C***. The latter matrix determines the stability of the species-density subsystem on its own with the juvenile fraction of each species equal to its equilibrium value. The matrix corresponds to the community matrix with elements ∂(*dC*_*i*_/*dt*)/∂*C*_*j*_ capturing the per-capita effect of the species in the community on each other’s growth rate. The community matrix determines the stability of a model, in which the dynamics of total species densities follow the same set of equations as in the full model, but the fraction of juveniles in the populations is constant over time and equal to the fraction of juveniles of the species at equilibrium (Figure 3, left-top panel). The comparison between the full Jacobian matrix and the community matrix reveals the impact of dynamic changes in the population structure of the species on the stability of the community equilibrium (see also Supplementary Information, section 2).

To assess the difference between constant and a dynamic juvenile fraction in the population, for all stable communities resulting from the stage-structured model with asymmetry in feeding and predation between juveniles and adults (*q* = 0.7, *ϕ* = 1.8) community dynamics were computed starting from the equilibrium community state using the reduced model including the differential equations *dC*_*i*_/*dt* for the species-density subsystem only, with the juvenile faction *Z*_*i*_ in each of the populations taken equal to its equilibrium value inferred from the stable community state (see Figure 3, right-top panel, Extended Data Figure 5 and Supplementary Information, section 2). Similarly, community dynamics were computed with the full model including the differential equations *dC*_*i*_/*dt* for the species-density subsystem and *dZ*_*i*_/*dt* for the species-structure subsystem starting from a community state in which the initial density of each species was exactly 50% of its equilibrium value inferred from the stable community state (see Figure 3, right-bottom panel and Extended Data Figure 5). For stable communities the extent of self-regulation of species is assessed with the diagonal elements of the community matrix, the *m*×*m* matrix ∂**K**/∂***C***, which measures the positive or negative effect of the total species abundance *C*_*i*_ on its own rate of change *dC*_*i*_/*dt* (see Extended Data Figure 4).

