## Supplementary information for "Dynamic population stage-structure stabilizes complex ecological communities"

### – Supporting Information –

André M. de Roos*

Institute for Biodiversity and Ecosystem Dynamics, University of Amsterdam, Amsterdam, The Netherlands

and

Santa Fe Institute, Santa Fe, New Mexico 87501, USA

May 1, 2020

### Supplementary Methods

Supplementary methods are presented in the following sections:

**1 Model simplification in case of ontogenetic symmetry** **2**

**2 Computing eigenvalues of the stage-structured model** **4**

**3 Stage-structured biomass model of species dynamics** **11**

*

#### 1 Model simplification in case of ontogenetic symmetry

The stage-structured model in terms of the juvenile and adult densities $J_i$ and $A_i$, respectively, can be reformulated into a model in terms of the total number of individuals of species $i$ and the fraction of juveniles in the population of species $i$. Define $C_i$ as the total density, $C_i = J_i + A_i$, and $Z_i$ as the fraction of juveniles of species $i$, $Z_i = J_i/C_i$. Using these alternative model variables the functional response value for the basal species can be written as:

$$F_1 = \frac{P}{\delta + \alpha_1 \left(qZ_1 + (2-q)(1-Z_1)\right) C_1} \tag{S1}$$

and the encounter rate of all non-basal species with their prey as

$$E_i = \sum_{k<i} \psi_{ik} \left(\phi Z_k + (2-\phi)(1-Z_k)\right) C_k \tag{S2}$$

From the ordinary differential equations (ODEs) for the juvenile and adult densities $J_i$ and $A_i$ presented in the methods section, the following system of ODEs for the alternative model variables $C_i$ and $Z_i$ can then be derived through analytical manipulation:

$$\begin{aligned}\frac{dC_i}{dt} &= b_i(F_i)(1-Z_i)C_i - \mu_i C_i \\ &\quad - \left(\phi Z_i + (2-\phi)(1-Z_i)\right) C_i \sum_{k>i} \alpha_k \psi_{ki} \frac{\left(qZ_k + (2-q)(1-Z_k)\right) C_k}{H_k + E_k}\end{aligned} \tag{S3}$$

$$\begin{aligned}\frac{dZ_i}{dt} &= b_i(F_i)(1-Z_i)^2 - m_i(F_i)Z_i \\ &\quad - 2(\phi-1)(1-Z_i)Z_i \sum_{k>i} \alpha_k \psi_{ki} \frac{\left(qZ_k + (2-q)(1-Z_k)\right) C_k}{H_k + E_k}\end{aligned} \tag{S4}$$

Assuming ontogenetic symmetry in ingestion and per-capita predation risk between juveniles and adults is equivalent to setting both $q$ and $\phi$ equal to 1, which simplifies the per-capita reproduction and maturation rate to:

$$b_i(F_i) = m_i(F_i) = \max\left(\gamma_i F_i - T_i, 0\right)$$

while the expressions for the encounter rate of non-basal species with their prey, $E_i$, equals:

$$E_i \;=\; \sum_{k<i} \psi_{ik} C_k \tag{S5}$$

The functional response for species $i$ is hence given by:

$$F_i = \begin{cases} \dfrac{P}{\delta + \alpha_1 C_1} & i = 1 \\[2ex] \dfrac{\sum_{k<i} \psi_{ik} C_k}{H_i + \sum_{k<i} \psi_{ik} C_k} & \text{otherwise} \end{cases} \tag{S6}$$

The equations describing the dynamics of total species densities $C_i$ and fractions of juveniles $Z_i$ therefore simplify to:

$$\frac{dC_i}{dt} = \max\left(\gamma_i F_i - T_i, 0\right)(1 - Z_i) C_i \;-\; \mu_i\, C_i \;-\; \sum_{k>i} \alpha_k \psi_{ki} \frac{C_k}{H_k + E_k}\, C_i \tag{S7}$$

$$\frac{dZ_i}{dt} = \max\left(\gamma_i F_i - T_i, 0\right)\left(1 - 3Z_i + Z_i^2\right) \tag{S8}$$

For all populations (basal and non-basal) the dynamics of the fraction of juveniles $Z_i$ hence follows a separable function, consisting of a factor $\max\left(\gamma_i F_i - T_i, 0\right)$ that only depends on the total species densities $C_i$ and a factor $\left(1 - 3Z_i + Z_i^2\right)$ that only depends on the fraction of juveniles $Z_i$. Irrespective of the fluctuations in the total species densities $C_i$, the fraction of juveniles will therefore approach the root of the quadratic condition $\left(1 - 3Z_i + Z_i^2\right) = 0$ for $t \to \infty$, i.e. approach the constant value:

$$\overline{Z} \;=\; \frac{3}{2} \,-\, \frac{1}{2}\sqrt{5} \;\approx\; 0.38 \tag{S9}$$

In the long run the dynamics of this juvenile-adult abundance model are therefore captured by a model that only considers total species abundances:

$$\frac{dC_i}{dt} = \max\left(\gamma_i F_i - T_i, 0\right)(1 - \overline{Z}) C_i \;-\; \mu_i\, C_i \;-\; \sum_{k>i} \alpha_k \psi_{ki} \frac{C_k}{H_k + E_k}\, C_i \tag{S10}$$

with $E_i$ and $F_i$ given by eq.(S5) and eq.(S6), respectively.

#### 2 Computing eigenvalues of the stage-structured model

To verify the local stability of the community states that appear to be stable based on numerical simulations I compute the eigenvalues characterising the dynamics in the neighbourhood of the equilibrium using the Jacobian matrix. In the neighbourhood of an equilibrium of the stage-structured model persistence of a species in the community guarantees that starvation does not occur for the juveniles nor the adults. In such a close neighbourhood of an equilibrium state the reproduction and maturation rate of adult and juvenile consumers are therefore necessarily positive, such that

$$b_i(F_i) = \max\left((2-q)\gamma_i F_i - T_i, 0\right) = (2-q)\gamma_i F_i - T_i \tag{S11}$$

$$m_i(F_i) = \max\left(q\gamma_i F_i - T_i, 0\right) = q\gamma_i F_i - T_i \tag{S12}$$

The dynamics of the total species densities $C_i$ and the fraction of juveniles in the populations $Z_i$ can then be described by simplified versions of the ODEs. (S3) and (S4):

$$\begin{aligned}\frac{dC_i}{dt} &= \left((2-q)\gamma_i F_i - T_i\right)(1-Z_i)C_i - \mu_i C_i \\ &\quad - \left(\phi Z_i + (2-\phi)(1-Z_i)\right) C_i \sum_{k>i} \alpha_k \psi_{ki} \frac{\left(qZ_k + (2-q)(1-Z_k)\right) C_k}{H_k + E_k}\end{aligned} \tag{S13}$$

$$\begin{aligned}\frac{dZ_i}{dt} &= \left((2-q)\gamma_i F_i - T_i\right)(1-Z_i)^2 - \left(q\gamma_i F_i - T_i\right) Z_i \\ &\quad - 2(\phi-1)(1-Z_i)Z_i \sum_{k>i} \alpha_k \psi_{ki} \frac{\left(qZ_k + (2-q)(1-Z_k)\right) C_k}{H_k + E_k}\end{aligned} \tag{S14}$$

The whole system of differential equations can be summarised as:

$$\frac{d\mathbf{C}}{dt} = \mathbf{K}\left(\mathbf{C}, \mathbf{Z}\right) \tag{S15}$$

$$\frac{d\mathbf{Z}}{dt} = \mathbf{L}\left(\mathbf{C}, \mathbf{Z}\right) \tag{S16}$$

in which $\mathbf{C}$ and $\mathbf{Z}$ are the vectors of total species abundances and fractions of juveniles in all populations, respectively. The vector-valued functions $\mathbf{K}(\mathbf{C}, \mathbf{Z})$ and $\mathbf{L}(\mathbf{C}, \mathbf{Z})$ contain the right-hand side of the ODEs (S13) and (S14) for $dC_i/dt$ and $dZ_i/dt$, respectively. For a community with $m$ species the Jacobian matrix of this model is a $2m \times 2m$ matrix $\mathbf{J}$

of the form:

$$\mathbf{J} \;=\; \begin{pmatrix} \dfrac{\partial \mathbf{K}}{\partial \mathbf{C}} & \dfrac{\partial \mathbf{K}}{\partial \mathbf{Z}} \\ \dfrac{\partial \mathbf{L}}{\partial \mathbf{C}} & \dfrac{\partial \mathbf{L}}{\partial \mathbf{Z}} \end{pmatrix} \;=\; \begin{pmatrix} \mathbf{V^1}+\mathbf{W^1} & \mathbf{V^2}+\mathbf{W^2} \\ & \\ \mathbf{V^3}+\mathbf{W^3} & \mathbf{V^4}+\mathbf{W^4} \end{pmatrix} \tag{S17}$$

$\mathbf{V^1}$, $\mathbf{V^2}$, $\mathbf{V^3}$ and $\mathbf{V^4}$ are 4 $m \times m$ matrices that capture the direct effects of two species in the community on each other, while $\mathbf{W^1}$, $\mathbf{W^2}$, $\mathbf{W^3}$ and $\mathbf{W^4}$ are 4 $m \times m$ matrices that capture the indirect effects between two species that operates through a third species. More specifically, indirect effects occur between species because changes in the total density $C_j$ and the fraction of juveniles $Z_j$ influence the encounter rate $E_k$ of a consumer species $k$, which in turn affects the predation rate of species $k$ on species $i$ (last summation terms in ODEs above). Indirect effects hence involve interactions between a predator species $k$ and two of its prey species with indices $i$ and $j$.

The elements of the matrices $\mathbf{V^1}$, $\mathbf{V^2}$, $\mathbf{V^3}$ and $\mathbf{V^4}$ are defined as:

$$V^1_{i,j} \;=\; \frac{d}{dC_j}\left(dC_i/dt\right),\; V^2_{i,j} \;=\; \frac{d}{dZ_j}\left(dC_i/dt\right),\; V^3_{i,j} \;=\; \frac{d}{dC_j}\left(dZ_i/dt\right),\; V^4_{i,j} \;=\; \frac{d}{dZ_j}\left(dZ_i/dt\right)$$

Notice however that the derivatives with respect to $C_j$ and $Z_j$ in these expressions are evaluated while ignoring the indirect effects that will be captured by the matrices $\mathbf{W^1}$, $\mathbf{W^2}$, $\mathbf{W^3}$ and $\mathbf{W^4}$, that is, while treating the quantities $E_k$ in the predation mortality terms (last summation terms in ODEs (S13) and (S14)) as constants.

The entries $V^1_{ij}$ are given by:

$$V^1_{ij} = \begin{cases} \left((2-q)\gamma_1 \dfrac{\delta F_1^2}{P} - T_1\right)(1-Z_1) - \mu_1 & \\ \quad - \left(\phi Z_1 + (2-\phi)(1-Z_1)\right) \displaystyle\sum_{k>1} \alpha_k \psi_{k1} \frac{\left(qZ_k + (2-q)(1-Z_k)\right) C_k}{H_k + E_k} & i=j=1 \\ \left((2-q)\gamma_i F_i - T_i\right)(1-Z_i) - \mu_i & \\ \quad - \left(\phi Z_i + (2-\phi)(1-Z_i)\right) \displaystyle\sum_{k>i} \alpha_k \psi_{ki} \frac{\left(qZ_k + (2-q)(1-Z_k)\right) C_k}{H_k + E_k} & i=j\neq 1 \\ -\alpha_j \psi_{ji} \dfrac{\left(qZ_j + (2-q)(1-Z_j)\right)}{H_j + E_j} \left(\phi Z_i + (2-\phi)(1-Z_i)\right) C_i & i<j \\ (2-q)\gamma_i \dfrac{H_i}{(H_i+E_i)^2} \psi_{ij} \left(\phi Z_j + (2-\phi)(1-Z_j)\right)(1-Z_i) C_i & i>j \end{cases}$$

(S18)

In the above expressions for the matrix elements $V^1_{ij}$ with $i > j$ I have used the identities

$$\frac{dF_i}{dC_j} = \frac{dF_i}{dE_i}\frac{dE_i}{dC_j} = \frac{H_i}{(H_i+E_i)^2}\frac{dE_i}{dC_j} = \frac{H_i}{(H_i+E_i)^2}\psi_{ij}\left(\phi Z_j + (2-\phi)(1-Z_j)\right)$$

and

$$\begin{aligned}\frac{d(F_1C_1)}{dC_1} &= \frac{d}{dC_1}\frac{PC_1}{\delta + \alpha_1\left(qZ_1 + (2-q)(1-Z_1)\right)C_1} \\ &= \frac{\delta P}{\left(\delta + \alpha_1\left(qZ_1 + (2-q)(1-Z_1)\right)C_1\right)^2} \;=\; \frac{\delta F_1^2}{P}\end{aligned}$$

In an equilibrium state all per-capita growth rates $(dC_i/dt)/C_i$ vanish such that the entries of the matrix $\mathbf{V^1}$ simplify to:

$$V^1_{ij} = \begin{cases} (2-q)\gamma_1\left(\frac{\delta F_1}{P} - 1\right)F_1(1-Z_1) & i=j=1 \\ 0 & i=j\neq 1 \\ -\alpha_j\psi_{ji}\frac{(qZ_j + (2-q)(1-Z_j))}{H_j+E_j}\left(\phi Z_i + (2-\phi)(1-Z_i)\right)C_i & i<j \\ (2-q)\gamma_i\frac{H_i}{(H_i+E_i)^2}\psi_{ij}\left(\phi Z_j + (2-\phi)(1-Z_j)\right)(1-Z_i)C_i & i>j \end{cases} \tag{S19}$$

The entries $V^2_{ij}$ are given by:

$$V^2_{ij} = \begin{cases} -\left((2-q)\gamma_1\frac{(\delta+\alpha_1 qC_1)F_1^2}{P} - T_1\right)C_1 & \\ \quad -2(\phi-1)C_1\sum_{k>1}\alpha_k\psi_{k1}\frac{(qZ_k+(2-q)(1-Z_k))C_k}{H_k+E_k} & i=j=1 \\ -\left((2-q)\gamma_iF_i - T_i\right)C_i & \\ \quad -2(\phi-1)C_i\sum_{k>i}\alpha_k\psi_{ki}\frac{(qZ_k+(2-q)(1-Z_k))C_k}{H_k+E_k} & i=j\neq 1 \\ -\alpha_j\psi_{ji}\frac{2(q-1)C_j}{H_j+E_j}\left(\phi Z_i + (2-\phi)(1-Z_i)\right)C_i & i<j \\ (2-q)\gamma_i\frac{H_i}{(H_i+E_i)^2}\psi_{ij}2(\phi-1)C_j(1-Z_i)C_i & i>j \end{cases} \tag{S20}$$

To derive the expressions for the matrix elements $V^2_{ij}$ with $i > j$ I have used the identities

$$\frac{dF_i}{dZ_j} = \frac{dF_i}{dE_i}\frac{dE_i}{dZ_j} = \frac{H_i}{(H_i+E_i)^2}\frac{dE_i}{dZ_j} = \frac{H_i}{(H_i+E_i)^2}\psi_{ij}2\left(\phi-1\right)C_j$$

and

$$\begin{aligned}\frac{d(F_1(1-Z_1))}{dZ_1} &= \frac{d}{dZ_1}\frac{P(1-Z_1)}{\delta+\alpha_1\left(qZ_1+(2-q)(1-Z_1)\right)C_1} \\ &= -\frac{\left(\delta+\alpha_1 qC_1\right)P}{\left(\delta+\alpha_1\left(qZ_1+(2-q)(1-Z_1)\right)C_1\right)^2} \\ &= -\frac{\left(\delta+\alpha_1 qC_1\right)F_1^2}{P}\end{aligned}$$

The entries $V^3_{ij}$ are given by:

$$V^3_{ij} = \begin{cases} -\gamma_1\dfrac{\alpha_1\left(qZ_1+(2-q)(1-Z_1)\right)F_1^2}{P}\left((2-q)(1-Z_1)^2-qZ_1\right) & i=j=1 \\ 0 & i=j\neq 1 \\ -\alpha_j\psi_{ji}\dfrac{\left(qZ_j+(2-q)(1-Z_j)\right)}{H_j+E_j}2(\phi-1)(1-Z_i)Z_i & i<j \\ \gamma_i\dfrac{H_i}{(H_i+E_i)^2}\psi_{ij}\left(\phi Z_j+(2-\phi)(1-Z_j)\right)\left((2-q)(1-Z_i)^2-qZ_i\right) & i>j \end{cases} \tag{S21}$$

To derive the expressions for the matrix elements $V^3_{ij}$ with $i = j = 1$ I have used the identity

$$\begin{aligned}\frac{dF_1}{dC_1} &= \frac{d}{dC_1}\frac{P}{\delta+\alpha_1\left(qZ_1+(2-q)(1-Z_1)\right)C_1} \\ &= -\frac{\alpha_1\left(qZ_1+(2-q)(1-Z_1)\right)P}{\left(\delta+\alpha_1\left(qZ_1+(2-q)(1-Z_1)\right)C_1\right)^2} \\ &= -\frac{\alpha_1\left(qZ_1+(2-q)(1-Z_1)\right)F_1^2}{P}\end{aligned}$$

Finally, the entries $V_{ij}^4$ are given by:

$$V_{ij}^4 = \begin{cases} -\gamma_1 \dfrac{2\alpha_1\,(q-1)\,C_1 F_1^2}{P}\big((2-q)(1-Z_1)^2 - qZ_1\big) \\ \quad -2\left((2-q)\gamma_1 F_1 - T_1\right)(1-Z_1) - (q\gamma_1 F_1 - T_1) \\ \quad -2(\phi-1)(1-2Z_1)\displaystyle\sum_{k>1}\alpha_k\psi_{k1}\frac{(qZ_k + (2-q)(1-Z_k))\,C_k}{H_k+E_k} & i=j=1 \\ \\ -2\left((2-q)\gamma_i F_i - T_i\right)(1-Z_i) - (q\gamma_i F_i - T_i) \\ \quad -2(\phi-1)(1-2Z_i)\displaystyle\sum_{k>i}\alpha_k\psi_{ki}\frac{(qZ_k + (2-q)(1-Z_k))\,C_k}{H_k+E_k} & i=j\neq 1 \\ \\ -\alpha_j\psi_{ji}\dfrac{2\,(q-1)\,C_j}{H_j+E_j}2(\phi-1)(1-Z_i)Z_i & i<j \\ \\ \gamma_i\dfrac{H_i\psi_{ij}2\,(\phi-1)\,C_j}{(H_i+E_i)^2}\big((2-q)(1-Z_i)^2 - qZ_i\big) & i>j \end{cases} \tag{S22}$$

The derivation of the expressions for the matrix elements $V_{ij}^4$ with $i = j = 1$ is based on the identity

$$\begin{aligned} \frac{dF_1}{dZ_1} &= \frac{d}{dZ_1}\frac{P}{\delta + \alpha_1\,(qZ_1 + (2-q)(1-Z_1))\,C_1} \\ &= -\frac{2\alpha_1\,(q-1)\,C_1 P}{\left(\delta + \alpha_1\,(qZ_1 + (2-q)(1-Z_1))\,C_1\right)^2} \\ &= -\frac{2\alpha_1\,(q-1)\,C_1 F_1^2}{P} \end{aligned}$$

As explained above, indirect effects occur between species because changes in the total density $C_j$ and fraction of juveniles $Z_j$ influence the encounter rate $E_k$ of a consumer species $k$, which in turn affects the predation rate of species $k$ on species $i$. These indirect effects therefore always arise because of the summation terms representing predation mortality in eqs. (S13) and (S14). In the predation rate of species $k$ only the term $1/(H_k + E_k)$ depends on the total density $C_j$ and the fraction of juveniles $Z_j$ of species

$j$ and the derivatives of this term with respect to $C_j$ and $Z_j$ equal

$$-\frac{1}{(H_k+E_k)^2}\psi_{kj}\left(\phi Z_j+(2-\phi)(1-Z_j)\right)$$

and

$$-\frac{1}{(H_k+E_k)^2}\psi_{kj}2\left(\phi-1\right)C_j$$

respectively. The elements of the matrices $\mathbf{W^1}$, $\mathbf{W^2}$, $\mathbf{W^3}$ and $\mathbf{W^4}$ are hence defined as:

$$\begin{aligned}
W^1_{i,j} &= \left(\phi Z_i+(2-\phi)(1-Z_i)\right)C_i\left(\phi Z_j+(2-\phi)(1-Z_j)\right)\\
&\quad\times\sum_{k>i}\alpha_k\psi_{ki}\psi_{kj}\frac{\left(qZ_k+(2-q)(1-Z_k)\right)C_k}{(H_k+E_k)^2}\\
W^2_{i,j} &= \left(\phi Z_i+(2-\phi)(1-Z_i)\right)C_i\;2(\phi-1)C_j\\
&\quad\times\sum_{k>i}\alpha_k\psi_{ki}\psi_{kj}\frac{\left(qZ_k+(2-q)(1-Z_k)\right)C_k}{(H_k+E_k)^2}\\
W^3_{i,j} &= 2(\phi-1)(1-Z_i)Z_i\left(\phi Z_j+(2-\phi)(1-Z_j)\right)\\
&\quad\times\sum_{k>i}\alpha_k\psi_{ki}\psi_{kj}\frac{\left(qZ_k+(2-q)(1-Z_k)\right)C_k}{(H_k+E_k)^2}\\
W^4_{i,j} &= 2(\phi-1)(1-Z_i)Z_i\;2(\phi-1)C_j\\
&\quad\times\sum_{k>i}\alpha_k\psi_{ki}\psi_{kj}\frac{\left(qZ_k+(2-q)(1-Z_k)\right)C_k}{(H_k+E_k)^2}
\end{aligned}$$

Note that $i$ and $j$ may be equal to each other as changes in the total density of $C_i$ and the fraction of juveniles $Z_i$ change the predation rate of species $k$ on species $i$ through a change in the functional response of species $k$, which effect is not captured by the matrices $\mathbf{V^1}$, $\mathbf{V^2}$, $\mathbf{V^3}$ and $\mathbf{V^4}$. Furthermore, note that all elements $W^1_{ij}$ are positive for species that are exposed to predation and equal to 0 only for top predators. Together with the fact that $V^1_{ii}=0$ for $i\neq 0$ this implies that the effect of species density $C_i$ on its own rate of change $dC_i/dt$ is 0 for top predators and positive for all non-basal species experiencing predation.

For all communities resulting from the stage-structured model that appear to be stable (defined as all communities for which the minimum and maximum values of the total species density differ less than $1.0 \cdot 10^{-6}$ from each other for all species) the Jacobian matrix is evaluated by substituting the average total abundance and fraction of juveniles for all species as well as all general and species-specific parameters into the matrices $\mathbf{V^1}$, $\mathbf{V^2}$, $\mathbf{V^3}$, $\mathbf{V^4}$, $\mathbf{W^1}$, $\mathbf{W^2}$, $\mathbf{W^3}$ and $\mathbf{W^4}$. The eigenvalues of the Jacobian matrix $\mathbf{J}$ (see equation (S17)) are subsequently computed numerically using the routine eigen() in R[1]. These calculations of the Jacobian matrix based on the analytical expressions for the matrices $\mathbf{V^1}$, $\mathbf{V^2}$, $\mathbf{V^3}$, $\mathbf{V^4}$, $\mathbf{W^1}$, $\mathbf{W^2}$, $\mathbf{W^3}$ and $\mathbf{W^4}$ were verified by also computing the Jacobian matrix numerically using central differencing methods applied to the right-hand side of the ODEs (S13) and (S14) for $dC_i/dt$ and $dZ_i/dt$.

To evaluate the effect of changes in the stage-composition of species (i.e. changes in the juvenile-adult ratio) on the stability of the community equilibrium, I compare the eigenvalues thus computed with the eigenvalues of a reduced model, in which for all species the fraction of juveniles in the population is assumed constant and equal to the fraction of juveniles of the species at equilibrium. More specifically, this reduced model is described by the ODE (S13) with the fraction of juveniles $Z_i$ set equal to its equilibrium value $\tilde{Z}_i$:

$$\begin{aligned}\frac{dC_i}{dt} &= \left((2-q)\gamma_i F_i - T_i\right)(1-\tilde{Z}_i)C_i + \mu_i C_i \\ &\quad - \sum_{k>i} \alpha_k \psi_{ki} \frac{\left(\phi\tilde{Z}_i + (2-\phi)(1-\tilde{Z}_i)\right) C_i}{H_k + E_k} \left(q\tilde{Z}_k + (2-q)(1-\tilde{Z}_k)\right) C_k \qquad \text{(S23)}\end{aligned}$$

Comparing the eigenvalues of this reduced model with the eigenvalues of the full model, in which the juvenile fraction in the population $Z_i$ is dynamic and changes at the same time scale as the total species density, reveals the impact of dynamic changes in the population structure of the species on the stability of the community equilibrium. The eigenvalues of the reduced model can be computed from its Jacobian matrix which equals the matrix $\partial\mathbf{K}/\partial\mathbf{C} = \mathbf{V^1} + \mathbf{W^1}$. The matrix $\partial\mathbf{K}/\partial\mathbf{C}$ captures the per-capita effect of species in the community on each other's growth rate and is hence equivalent to the community interaction matrix that determines stability in community models without population structure.

### 3 Stage-structured biomass model of species dynamics

To check the robustness of the results obtained with the stage-structured model in terms of juvenile and adult numerical densities, numerical simulations of community dynamics were also carried out, using a stage-structured biomass model for species dynamics[2]. More specifically, each species was represented by 3 life history stages, referred to as juveniles, subadults and adults. Such a stage-structured biomass model[2] constitutes an approximation to a size-structured population model that accounts for a complete size distribution of individuals between their size at birth and size at maturation, in which the rates of feeding, metabolic maintenance, somatic growth, and reproduction all scale linearly with individual body size[3]. Juvenile and subadult individuals are assumed to use their net-energy production (the difference between assimilation and metabolic maintenance rate) for somatic growth, whereas adults are assumed not to grow and use their net-energy production for reproduction. Dynamics are in terms of juvenile, subadult and adult biomass densities, indicated with $J_i$, $S_i$ and $A_i$, respectively.

In the absence of predation the life history processes in the stage-structured biomass model are described by the following mass-specific rate functions:

$$\text{Juvenile somatic growth} \qquad g_i^J(F_i) = \max\left((2-q)\gamma_i F_i - T_i, 0\right) \tag{S24}$$

$$\text{Subadult somatic growth} \qquad g_i^S(F_i) = \max\left(q\gamma_i F_i - T_i, 0\right) \tag{S25}$$

$$\text{Adult reproduction} \qquad b_i(F_i) = \max\left((2-q)\gamma_i F_i - T_i, 0\right) \tag{S26}$$

$$\text{Juvenile mortality} \qquad d_i^J(F_i) = \mu_i - \min\left((2-q)\gamma_i F_i - T_i, 0\right) \tag{S27}$$

$$\text{Subadult mortality} \qquad d_i^S(F_i) = \mu_i - \min\left(q\gamma_i F_i - T_i, 0\right) \tag{S28}$$

$$\text{Adult mortality} \qquad d_i^A(F_i) = \mu_i - \min\left((2-q)\gamma_i F_i - T_i, 0\right) \tag{S29}$$

$$\text{Juvenile maturation} \qquad m_i^J(F_i) = \begin{cases} \dfrac{g_i^J(F_i) - D_i^J}{1 - z^{1-D_i^J/g_i^J(F_i)}} & \text{if } g_i^J(F_i) > 0 \\ 0 & \text{otherwise} \end{cases} \tag{S30}$$

$$\text{Subadult maturation} \qquad m_i^S(F_i) = \begin{cases} \dfrac{g_i^S(F_i) - D_i^J}{1 - z^{1-D_i^S/g_i^S(F_i)}} & \text{if } g_i^S(F_i) > 0 \\ 0 & \text{otherwise} \end{cases} \tag{S31}$$

In these equations $F_i$ represents the functional response of species $i$, which for the basal species equals:

$$F_1 \;=\; \frac{P}{\delta \,+\, \alpha_1\left(\left(2-q\right)J_1 + q\,S_1 + \left(2-q\right)A_1\right)} \tag{S32}$$

The parameter $q$ in this expression determines the asymmetry in feeding capacity between juveniles, subadults and adults (for the purpose of this study taken the same for all species). Non-basal species forage following a type II functional response:

$$F_i \;=\; \frac{E_i}{H_i + E_i} \tag{S33}$$

in which $E_i$ represents the encounter rate of non-basal species $i$ with prey biomass:

$$E_i \;=\; \sum_{k<i} \psi_{ik}\left(\phi\,J_k + \left(2-\phi\right)S_k + \left(2-\phi\right)A_k\right) \tag{S34}$$

The parameter $\phi$ represents the bias of the consumer species toward feeding on juvenile as opposed to subadult and adult prey (for the purpose of this study taken the same for all species). Notice that all species are ordered according to their body size and hence species $i$ can only feed on species with an index $k < i$.

The parameter $T_i$ in the life history functions (S24)-(S31) represents the (mass-specific) loss rate through metabolic maintenance requirements, while the parameter $\mu_i$ represents the background mortality. The parameter $\gamma_i$ determines the maximum assimilation rate per unit biomass, while the parameter $z$ equals the ratio between the body size at entering and leaving each of the immature stages (the juvenile and subadult stage). The parameters $\gamma_i$, $q$ and $T_i$ also determine the minimum food availability that is needed by juveniles, subadults and adults to just keep itself alive without producing any offspring and without maturing.

$D_i^J$ and $D_i^S$ indicate the total mortality rate experienced by juvenile and subadult individuals, respectively, which in the absence of predation equals $\mu_i$, but in the presence of predation also includes the predation mortality (see below; note that $D_i^J$ and $D_i^S$ do not include starvation mortality as starvation mortality only occurs when $g_i^J(F_i) = 0$ or $g_i^S(F_i) = 0$, in which case $m_i^J(F_i) = 0$ and $m_i^S(F_i) = 0$, respectively). Equations (S26), (S24), (S25), (S30) and (S31) express that adult reproduction, juvenile and subadult growth in body size and juvenile and subadult maturation come to a halt under starvation conditions, which for juveniles, subadults and adults occur when $q\gamma_i F_i < T_i$ and $(2-q)\gamma_i F_i < T_i$, respectively. Under these starvation conditions juveniles, subadults and adults experience increased mortality (eqs.(S27), (S28) and (S29)). The mass-specific juvenile and subadult maturation rates depends on both juvenile and subadult growth rate in body size as well as total juvenile and subadult mortality, $D_i^J$ and $D_i^S$, respectively. The functional form of the maturation rates $m_i^J(F_i)$ and $m_i^S(F_i)$ are chosen such that any equilibrium state predicted by the stage-structured biomass model corresponds uniquely to an equilibrium state of a structured model that accounts for a complete size distribution of individuals between their size at birth and size at maturation, in which the rates of feeding, metabolic maintenance, somatic growth, and reproduction all scale linearly with individual body size[3].

A representation of each species by 3 life history stages with the smallest juveniles most vulnerable to predation mortality and the maturation of the larger immature individuals most limited by food availability was chosen because the dynamics of such a 3-stage biomass model has been found to closely resemble the dynamics of population models with a complete size distribution that are based on a dynamic energy budget model for the individual energetics[4]. Similar results as presented in Extended Data Figure 6, 7 and 8 have, however, also been obtained using a stage-structured biomass model with only a single juvenile and adult life history stage to describe species dynamics (results not shown).

The dynamics of the juvenile, subadult and adult biomass densities of all species in the community are now described by the following set of ordinary differential equations (ODEs):

$$\begin{aligned}\frac{dJ_i}{dt} &= b_i(F_i)A_i + g_i^J(F_i)J_i - m_i^J(F_i)J_i - d_i^J(F_i)J_i \\ &\quad - \phi J_i \sum_{k>i} \alpha_k \psi_{ki} \frac{(2-q)J_k + qS_k + (2-q)A_k}{H_k + E_k}\end{aligned} \tag{S35}$$

$$\begin{aligned}\frac{dS_i}{dt} &= m_i^J(F_i)J_i + g_i^S(F_i)S_i - m_i^S(F_i)S_i - d_i^S(F_i)J_i \\ &\quad - (2-\phi) S_i \sum_{k>i} \alpha_k \psi_{ki} \frac{(2-q)J_k + qS_k + (2-q)A_k}{H_k + E_k}\end{aligned} \tag{S36}$$

$$\begin{aligned}\frac{dA_i}{dt} &= m_i^S(F_i)S_i - d_i^A(F_i)A_i \\ &\quad - (2-\phi) A_i \sum_{k>i} \alpha_k \psi_{ki} \frac{(2-q)J_k + qS_k + (2-q)A_k}{H_k + E_k}\end{aligned} \tag{S37}$$

In these equations $E_k$ and $H_k$ represent the encounter rate with prey and the half-saturation density in the functional response of species $k$, respectively (eq.(S34)), while the parameters $q$ and $\phi$ represent the asymmetry in foraging rate and predation risk, respectively, between juvenile, subadult and adult individuals. The default values for these parameters equal 1, implying that all 3 stages have identical life history rates ($q = 1$) and that consumers feed indiscriminately on juveniles, subadults and adults of their prey species ($\phi = 1$). Finally, the parameter $\alpha_k$ represents the maximum (mass-specific) foraging rate of consumer species $k$.

Given the above equations, the total juvenile mortality rate, on which the maturation rate (eq.(S30)) of juvenile into subadult biomass depends, is the sum of background (but not starvation) mortality and predation mortality:

$$D_i^J = \mu_i + \phi \sum_{k>i} \alpha_k \psi_{ki} \frac{(2-q)J_k + qS_k + (2-q)A_k}{H_k + E_k} \tag{S38}$$

Starvation mortality is excluded from $D_i^J$ because the maturation rate equals 0 under starvation conditions. Analogously, the total subadult mortality rate, on which the maturation rate (eq.(S31)) of subadult into adult biomass depends, is the sum of background (but not starvation) mortality and predation mortality:

$$D_i^S = \mu_i + \phi \sum_{k>i} \alpha_k \psi_{ki} \frac{(2-q)J_k + qS_k + (2-q)A_k}{H_k + E_k} \tag{S39}$$

#### Model parameterisation and numerical simulation details

Parameterisation of the stage-structured biomass model follows the same procedure as the stage-structured model in terms of juvenile and adult abundance (see Methods). In short, half-saturation prey densities $H_i$ for non-basal species are sampled from a uniform distribution on the interval $[0.5, 2.5]$. The ratio between the smallest and the largest body size in each of the two immature life stages, $z$, that occurs in the maturation rates of the stage-structured biomass model (eq.(S30) and (S31)), is for all species taken the same and equal to $z = 0.1$. Individuals therefore are hence assumed to grow 2 orders of magnitude in body size between birth and maturation. The parameters $\alpha_i$, $\gamma_i$, $T_i$ and $\mu_i$ all represent (mass-specific) rates and are assumed to scale with $w_i^{-0.25}$ following the equations:

$$\begin{aligned}
\alpha_i = & \ \ \alpha_0 \left(1 + 2\sigma_\alpha \left(x_{i1} - \tfrac{1}{2}\right)\right) w_i^{-0.25} \\
\gamma_i = & \ \ \gamma_0 \left(1 + 2\sigma_\gamma \left(x_{i2} - \tfrac{1}{2}\right)\right) w_i^{-0.25} \\
T_i = & \ \ T_0 \left(1 + 2\sigma_T \left(x_{i3} - \tfrac{1}{2}\right)\right) w_i^{-0.25} \\
\mu_i = & \ \ \mu_0 \left(1 + 2\sigma_\mu \left(x_{i4} - \tfrac{1}{2}\right)\right) w_i^{-0.25}
\end{aligned} \tag{S40}$$

The default mean values of the species-specific parameters equal $\alpha_0 = 1.0$, $\gamma_0 = 0.6$, $T_0 = 0.1$ and $\mu_0 = 0.015$[4], while the species-specific parameters $\alpha_i$, $\gamma_i$, $T_i$ and $\mu_i$ are randomly selected from a Bates distribution of degree 3 around these mean values. A Bates distribution is the continuous probability distribution of the mean, $X$, of 3 independent uniformly distributed random variables on the unit interval. Random values from this distribution range between 0 and 1 with mean value of $\frac{1}{2}$ and are easily generated by taking the mean of 3 independent samplings from a uniform distribution on the unit interval $[0, 1]$. The quantities $x_{ij}$ are independent realisations of the random variable $X$, while $\sigma_\alpha$, $\sigma_\gamma$, $\sigma_T$ and $\sigma_\mu$ represent the one-sided, relative width of the distributions of the species-specific parameters $\alpha_i$, $\gamma_i$, $T_i$ and $\mu_i$, respectively, around the mean values $\alpha_0 = 1.0$, $\gamma_0 = 0.6$, $T_0 = 0.1$ and $\mu_0 = 0.015$. Default values for these relative widths equal 0.1, such that all species-specific parameters $\alpha_i$, $\gamma_i$, $T_i$ and $\mu_i$ range between 0.9 and 1.1 times their default mean value and follow hump-shaped distributions within these ranges. The productivity $P$ and turn-over rate $\delta$ of the exclusive resource of the basal species are taken equal to 60 and 2.0, respectively, in all computations. The two remaining parameters, the foraging asymmetry parameter $q$ and the predation asymmetry parameter $\phi$, in the model are varied between the different computations to assess their effect on community dynamics.

As described in the Methods section food webs are generated by selecting $N = 500$ random niche values $n_i$ uniformly from the interval $[0, 1]$ and associated with species body mass following $w_i = w_{max}^{n_i}/w_{min}^{1-n_i}$. Subsequently, the network of feeding interactions between these $N = 500$ species is constructed by generating for each non-basal species the midpoint of its feeding niche $c_i$ following the procedure and default values for the mean prey-predator body mass ratio described in the Methods and Extended Data Figure 9. Numerical integrations of the food web with $N = 500$ species are carried out using an adaptive Runge-Kutta (Cash-Karp) method implemented in C. Relative and absolute tolerances during the integration equal $1.0 \cdot 10^{-7}$ and $1.0 \cdot 10^{-13}$, respectively. During the first $10^4$ time units no species are removed from the community, even if they attain very low density. For $t > 10^4$ each species, whose total biomass density $J_i + A_i$ drops below $10^{-8}$, is removed from the community. This persistence threshold ensures that the product of the relative tolerance ($10^{-7}$) and the lowest species density ($10^{-8}$) is larger than the machine precision (equal to $1.11 \cdot 10^{-16}$ according to the IEEE 754-2008 standard in case of double precision). During numerical computations mean and variance as well as the maximum and minimum value of the total species biomass $J_i + A_i$ are continuously monitored for all species. The values of these measured statistics are reset whenever the community structure changes as one or more species in the community go extinct. Numerical integrations are halted whenever the community structure has not changed for $10^6$ time units and no change has occurred from one time unit to the next in the values of these statistics (mean, minimum, maximum and variance of total species density) for all species in the community.

4. De Roos, A. M. & Persson, L. *Population and community ecology of ontogenetic development* (Princeton University Press, Princeton, 2013).
